# Structural and Functional Versatility of the Amyloidogenic Antimicrobial Peptide Citropin 1.3

**DOI:** 10.1101/2025.01.31.635854

**Authors:** Fabio Strati, Mariana Pigozzi Cali, Yehudi Bloch, Siavash Mostafavi, Jim Monistrol, Aleksandr Golubev, Bader Rayan, Emil Gustavsson, Meytal Landau

## Abstract

Citropin 1.3 is an antimicrobial peptide produced by the amphibian *Litoria citropa* (Southern bell frog), which self-aggregates into distinct fibrillar structures, however, the function of the fibrils remains unclear and largely unexplored. In this study, the structural and functional properties of citropin 1.3 were investigated using cryogenic electron microscopy and fluorescence microscopy in the presence of membrane and cell models, and with X-ray crystallography. Canonical amyloids, multilayered nanotubes, and a novel mixed fibril were observed. Experiments with negatively charged giant unilamellar vesicles revealed that the peptide facilitates membrane fusion while simultaneously undergoing phase separation in the presence of phospholipids. In presence of mammalian cells, citropin 1.3 permeabilizes membranes, leading to cell death, and over time, colocalizes with genetic material. Overall, this work provides new insights into the structural dynamics of the amyloidogenic antimicrobial peptide citropin 1.3 and its interactions with different systems.

## Introduction

Protein and peptide self-assembly into amyloid fibrils is associated with a wide range of functions and applications. Of note, more than 20 human pathologies, including Alzheimer’s and Parkinson’s diseases, are amyloid-associated diseases whose hallmark is the presence of extracellular or intracellular deposits of insoluble protein aggregates defined as amyloid plaques ^1,2^. Self-assembly into amyloid fibrils is also connected to functional roles in the life cycles of different organisms. In bacteria, amyloid fibrils have been observed to act as biofilm stabilisers ^3,4^ and to serve as toxins for cell invasion ^4,5^. Some eukaryotic organisms secrete antimicrobial peptides (AMPs) that have been demonstrated to self-aggregate into amyloid fibrils which then attack and lyse pathogenic bacteria ^6^. Human amyloids linked to neurodegenerative and systemic diseases have also shown to exhibit antimicrobial activity, highlighting the significance of the amyloid-antimicrobial connection ^6–11^ and the importance of the investigation of novel AMPs.

Amyloids mostly share a typical cross-β structure formed by tightly packed β-sheets usually arranged in the so-called steric zipper conformation, an extremely stable motif responsible for the peculiar stiffness and resistance of amyloid fibrils. In the past years, high resolution studies of virulent and AMP fibrils revealed new morphologies extending beyond the canonical amyloid cross-β structure including the discovery of cross-α fibrils composed of stacked α-helices ^12,13^. Surprisingly, some of these peptides have been observed in biophysical and structural studies to form either or both cross-α and -β fibrils, a type of polymorphism hypothesized to be related to the functional role of the peptide itself.

The amphibian AMP uperin 3.5 is currently the only AMP known to exhibit both cross-α and cross- β fibril structures at high resolution, determined by X-ray crystallography and cryo-electron microscopy (cryo-EM), respectively ^6,14^. These structures revealed secondary structure polymorphism, which is highly unusual among proteins and peptides with available three-dimensional structures ^15^. Another amphibian AMP, aurein 3.3, forms amyloid fibrils with a cryo-EM structure that revealed an atypical fibril architecture composed of kinked β-sheets^14^. This structure resembles functional amyloids known as LARKS (low-complexity aromatic-rich kinked segments) ^16,17^. The fibril-forming potential of aurein 3.3 was predicted using a computational platform designed to identify fibril-forming AMPs ^15^. Using the same platform, citropin 1.3 was also identified as a potential fibril-forming peptide. Here, we report its structure and activity, providing further insights into the fibril-forming tendencies of AMPs.

Citropins are a family of peptides extensively studied for their broad-spectrum antibacterial activity against Gram positive and Gram negative bacteria ^18–20^, as well as their potential anticancer properties ^21,22^. In this study, we investigated the structure-activity relationship (SAR) of citropin 1.3 using various membrane models and mammalian/bacterial cells. Employing cryo-EM and X-ray crystallography, we determined its high-resolution structures, while fluorescence microscopy was used to observe dynamic cellular processes in real-time. Our analysis revealed a wide spectrum of fibrillar structures, ranging from canonical amyloid fibrils to nanotubes, demonstrating significant structural polymorphism influenced by the aqueous environment and the presence of lipid vesicles. Notably, fluorescence microscopy unveiled liquid-liquid phase separation (LLPS) of citropin 1.3 in the presence of phospholipids. In both bacterial and mammalian cellular environments, citropin 1.3 was internalized. In mammalian cells, it colocalized with genetic material and formed droplet-like condensates in the cytosol.

This work uncovers novel structural insights and extensive polymorphism at both the secondary and quaternary levels, offering a deeper understanding of the SAR, cellular interactions, and phase separation properties of citropin 1.3. These findings provide compelling evidence supporting the antimicrobial-amyloid link at high resolution, and particularly in the context of cellular toxicity. Moreover, this study introduces promising research directions to investigate the mechanisms underlying amyloid toxicity and explore their potential implications for future therapeutic applications.

## Results

### Condition-dependent aggregation of citropin 1.3 revealed by cryo-EM

Lyophilised citropin 1.3 was dissolved in different aqueous solutions to monitor its aggregation under different conditions. After following fibril growth via negative stain transmission electron microscopy (TEM), the samples were cryo-plunged and imaged. A total of three datasets were acquired in different conditions: I) 1 mM citropin 1.3 in 50 mM NaCl at pH 5; II) 1 mM citropin 1.3 in 10 mM phosphate-buffered saline (PBS) at pH 7.4; III) 500 μM citropin 1.3 in 10 mM PBS at pH 7.4 in the presence of 2.5mM negatively charged liposomes (composition is described in the Method section). The peptide was incubated for 48 h and 24 h in the datasets with and without liposomes respectively.

A distinct difference in aggregate morphology was observed, with canonical amyloids formed at pH 5 (Fig. S1A) while wide nanotubes formed at pH 7.4 (Fig. S1B). The presence of negatively charged liposomes induced the formation of three distinct canonical amyloids polymorphs (Fig. S1C). Unfortunately, no high-resolution structures have been obtained at pH 7.4 without liposomes since the formed nanotubes possessed multiple internal symmetries (Fig. S2) which made the data processing inconclusive. Therefore, this dataset will not be further discussed.

### Cryo-EM structures of citropin 1.3 at pH 5

Four main fibril morphologies have been identified and labelled from I to IV (Fig. S3A) according to their relative particle count after extraction. The presence in the corresponding 2D classes of layer lines perpendicular to the filament axis at a spacing of approximately 4.75 Å ^23,24^, together with the analysis of the averaged power spectra, clearly showed that three polymorphs (Pol II, III and IV) consisted of canonical amyloids with stacking of β-sheets along the z-axis (Fig. S3A).

We determined the 3D structure of Pol II and III fibrils, however we were not able to improve the resolution of Pol IV to sufficiently determine the staggering structure of β-strands. Nevertheless, the three polymorphs showed very distinct peptide arrangements indicating no transition of one polymorph to the other within the same fibril (Fig. S3A). The reconstructed volume of Pol II led to a map with 2.8 Å resolution displaying a C3i pseudo-symmetry, arranged in a triskeles motif, with three chains per layer (Fig. 1A). Due to reasons that might be connected to intrinsic chain flexibility, only the internal strands of the map could be solved at high enough resolution allowing us to model this part of the map, while it is possible that additional peptide chains flank the fibril core. The model showed excellent fitting of the full-length peptide into the obtained map (Fig. 1A). Each strand presented a kink of almost 90° at residue Lys8, which forms a *cis*-like bond with Lys7, displaying angles typical of a left-handed α-helix (Fig. S4E). The kink creates a motif similar to the one of the LARKS ^16,17^ and disruption of the β-sheets backbone hydrogen bonds (H-bonds) (Fig. S4A). H-bonds were formed between the positively charged amine of Lys8 and the carboxyl of C-terminal Leu16 from adjacent chains (Fig. 1A).

**Figure 1.**
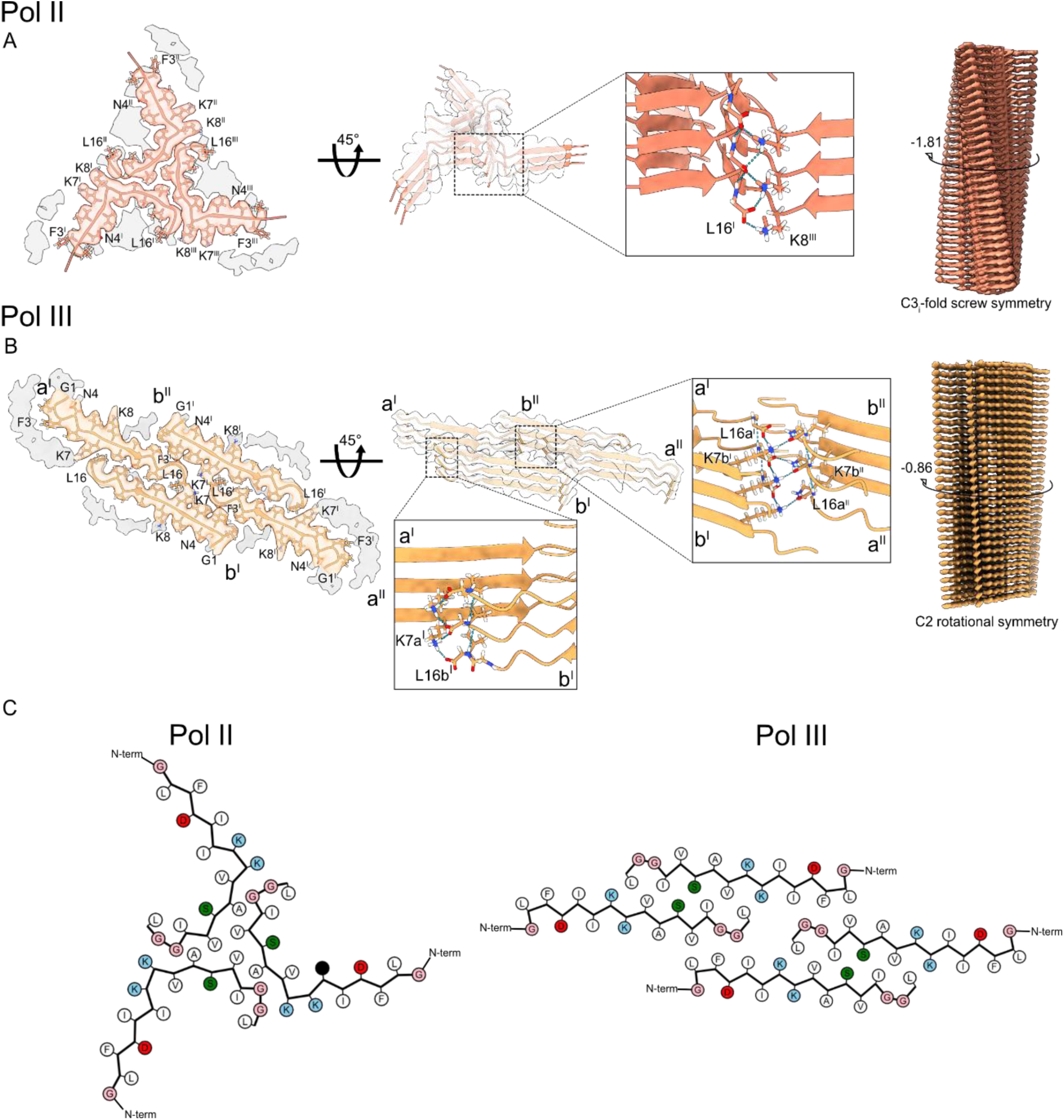
Amyloid structures of citropin 1.3 at pH 5. **A)** Polymorph II displaying three peptide chain per layer, marked ^I,II,III^. **B)** Polymorph III displaying four peptide chain per layer, marked a^I,^a^II^,b^I^,b^II^. Both polymorphs contain potential additional protein chains suggested by an incomplete map flanking the resolved fibril part. **A-B)** Left panels: atomic model of a single layer built into the Coulomb density with visualization of low-resolution densities. Middle panels: Three-layered density map featuring the fit of the atomic model, accompanied by close-up insets highlighting the formed hydrogen bond between the C-terminal carboxyl backbone group of Leu16^I^ and the side chain of Lys8^III^ (**A**) or between Leu16(a^I)^ and Lys7(b^I^), and Leu16(a^II)^ and Lys7(b^II^) (**B**). Left panels: full maps with reported twist. **C)** Cartoon of residue properties within the fibril cross-section. Hydrophobic, polar, and negatively/positively charged residues are indicated in white, green, red, and blue, respectively.

The structure of Pol III showed a C2 rotational symmetry and resolution of 3.1 Å with fully extended β-sheets and four monomers per layer, displaying extra peptide chain densities flanking the fibril core (Figure 1B). Strands **a^I^** and **b^I^**presented fully extended chains with the entire sequence fitting inside the cryo-EM density (Fig. 1B). Continuous β-sheets are formed for chain **a^I^** between Ile6 and Ser11, and for chain **b^I^** between Phe3 and Ser11. In addition to backbone H-bonds along the β-sheets, an interchain H-bond connects the N-terminal amine group of Gly1 and the carboxyl of Asp4 formed in each strand, potentially creating a turn at the N-terminal end (Figs. 1B,C & S5A). Finally, interchain electrostatic interactions potentially form between the amine of Lys7 side chain and the C-terminal carboxyl group of Leu16 in both chain pairs, resulting in a total of four electrostatic bonds per layer. Two of these bonds are facing the middle of the fibrillar arrangement. The two others are located at the outer surface of the fibril from both sides, which are potentially flanked by additional protein chains suggested by an incomplete and disordered map (Fig. 1B).

Finally, Pol I, as observable from the power spectra and from the 2D class averages (Figs. 2A and S6A) presented a novel fibril morphology. Reflections in different layer lines typical of helical polymers were present simultaneously to four reflections at 4.75 Å rotated by 52° from the meridian, and a ring at 10 Å, features that suggest the formation of an amyloid fibrils ^25–27^. However, the β-sheets do not stack perpendicular to the fibril axis as in most known amyloids, but at a 52° angle. An analysis of the fibrils in the micrographs (Fig. 2B), showed a crossover typical of amyloids, a feature also observed in the 2D class (Fig. S6A). 3D helical refinements without symmetry imposition performed with the software package cryoSPARC ^28^ generated a map with a resolution of 4.85 Å (Fig. S6B), revealing the formation of a nanotube with an oval cross-section made of three concentric layers (Fig. 2C,E). The obtained resolution did not allow the modelling of the peptide into the map densities. However, looking at each individual layer combined with the information from the power spectra, allowed us to speculate on its secondary structure arrangement. In the first, outer, layer, superimposing α-helices of citropin 1.3 predicted by AlphaFold (AlphaFoldDB: P81846) onto the map suggested the formation of an array of helices which surrounds the outer layer (Fig. 2C). In the second, middle, layer, we hypothesize that the densities consist of stacked β-sheets along the vertical axes with a tilt of 52° in agreement with the power spectra (Fig. 2A,C) and a measured distance between β-sheets of circa 5 Å (Figure 2D), suggesting that this layer might consist of amyloid-like β-sheets that do not stack perpendicular to the nanotube. The inner layer is too poorly resolved. Overall, this fibril polymorph showed a novel morphology which might represent the first case of a mixed amyloid-polymer fibril with both α- and β- secondary structures.

**Figure 2.**
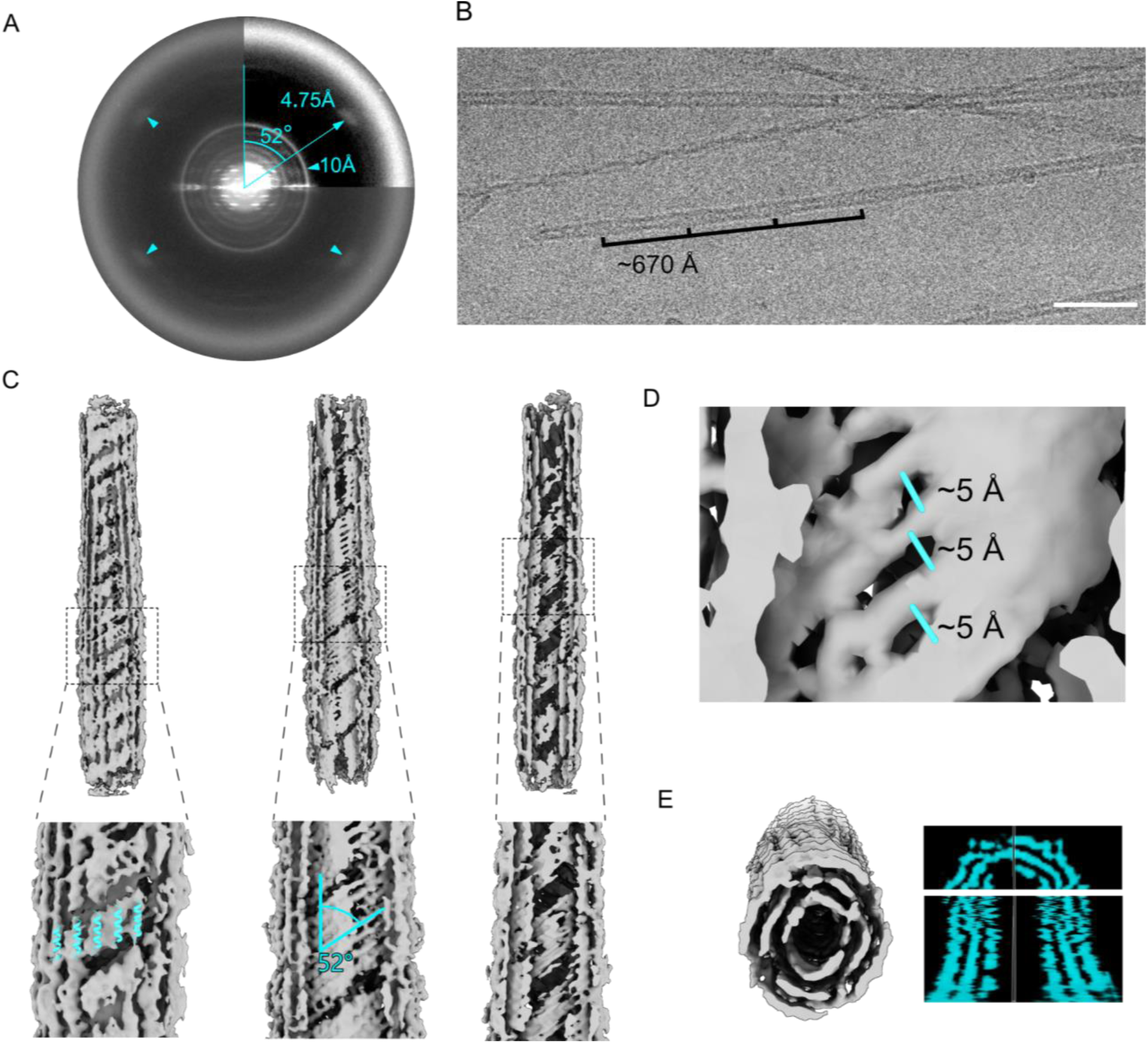
Polymorph I power spectra interpretation and map analysis. **A)** Averaged power spectra of Pol I, main visible features are the 4.75 Å peaks highlighted by triangular markers rotated by 52° from the meridian axes indicating stacking of β-sheets, and a 10 Å ring typical for lateral interaction of β-sheets, both common features of amyloid fibrils. However, the β-sheets to the fibril axis with at a 52° angle. **B)** Representative NaCl 50 mM pH 5 dataset micrograph with Pol I filament and visualisation of cross-over distance with 670 Å periodicity, scale bar 500 Å. **C)** Map of the reconstructed density with insights in each internal layer of the fibril. In the outer layer, predicted α-helices of the monomer (AlphaFoldDB: P81846) wrap the structures along the vertical axe, in the second, middle, layer, the formed densities resemble amyloid-like β-sheets, assumption supported by the 52° angle formed by the densities as well reported in power spectra. Finally, the third and most internal layer shows unresolved features. **D)** Measured distance between β-sheets of the middle layer reported to be about 5 Å. **E)** Cross-section and orthogonal view of the map with clear separation of three layers along the vertical (top) and horizontal (bottom) axes.

### Crystal structure of citropin 1.3 reveals α-helices assembling into a nanotube-like supramolecular structure

Citropin 1.3 was found to form cubic microcrystals, which diffracted to a resolution of 1.6 Å. The crystal structure revealed two chains of the 16-residue peptide within the asymmetric unit (Fig. S7A). These peptide chains adopt α-helical conformations, tightly packed through hydrophobic interactions involving residues such as Ile5, Val9, Val12, and Leu16. Additionally, a hydrogen bond is formed between Lys8 on chain B and Ser11 on chain A. Each chain also exhibits an internal electrostatic interaction between Lys8 and Asp4.

The α-helical dimers further assemble into a spiraling supramolecular structure (Fig. 3). Inter-dimer interactions include tight packing between Phe3 and Lys7 of neighboring chain Bs, alongside interactions with Leu16 from chain A (Fig. S7B). Furthermore, Ala10 and Ile13 from chains A and B of adjacent dimers engage in hydrophobic packing, as do Leu2 and Ile5 from neighboring A chains (Fig. S7C). These tightly packed helices collectively form a supramolecular assembly resembling a nanotube (Fig. 3). The fibril’s surface features alternating hydrophobic and positively charged zigzagged belts (Fig. 3). This unique surface topology likely facilitates interactions with and subsequent disruption of negatively charged lipid bilayers, such as bacterial membranes. The potential formation of such nanotubes in non-crystalline environments remains unexplored. Nonetheless, the structure confirms that citropin 1.3 can adopt α-helical conformations in addition to the β-strand-rich structures observed in cryo-EM studies. The crystal packing suggests a propensity for supramolecular assembly in the helical form, although its physiological relevance is yet to be determined.

**Figure 3.**
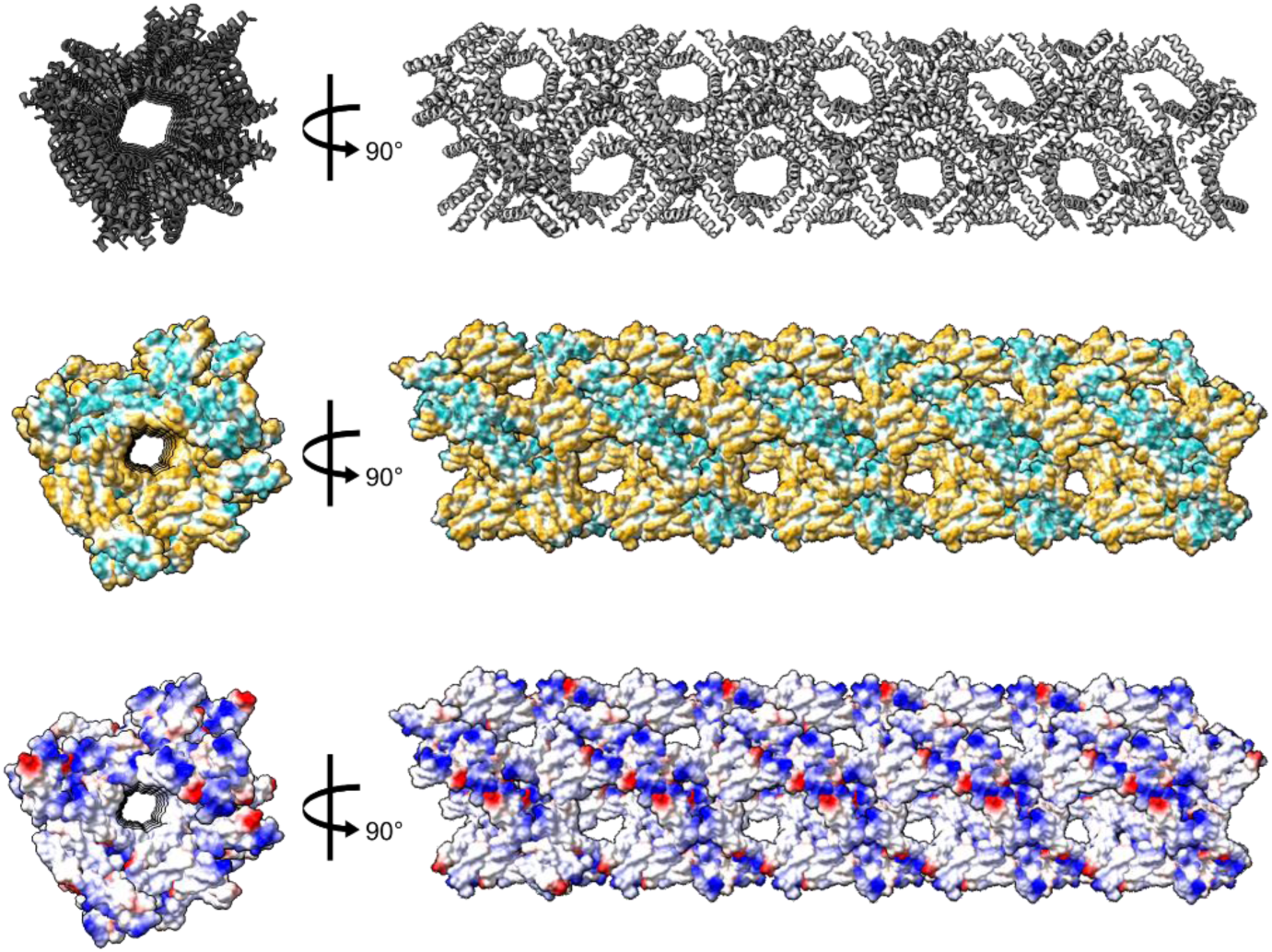
Crystal Structure of citropin 1.3 forms spiraling α-helices assembling into nanotubes. The figure illustrates the nanotube structure of citropin 1.3 from two perpendicular orientations, presented in three different representations: secondary structure with helices depicted in grey (top), and surface representations colored by hydrophobicity (middle pane) and electrostatics (bottom). The nanotube features a hydrophobic surface channel running along its length, with an approximate diameter of 40–50 Å. Due to the cubic nature of the crystal packing, these nanotubes are oriented along all dimensions of the crystal lattice.

### Cryo-EM structures of citropin 1.3 in presence of liposomes

To investigate the interactions and effects of citropin 1.3 on bacterial membranes, we employed a model system consisting of simple negatively charged liposomes, composed of multilamellar vesicles (details provided in the Methods section). Citropin 1.3 was incubated in PBS at pH 7.4 for several hours with the liposomes at a 1:5 molar ratio (peptide:liposome) and subsequently immobilized on cryo-EM grids. The resulting dataset revealed three distinct fibril morphologies (Fig. S8), which were resolved at high resolution.

The fibril with most particles, Pol I-L, presented a C3i screw symmetry with propeller shape (Fig. 4A) at a resolution of 2.98 Å showing three peptide chains per layer and a flexible region loosely bound to the main chain (Fig. S8). Pol I-L closely resembled the previous Pol II triskeles (Fig. 1A) in terms of symmetry and peptide arrangement, with some major differences in terms of β-sheets configuration and H-bonds formation. The main difference between the two fibrils was the extent of the kink observed around Lys8 in Pol II. In Pol I-L, instead of the kink there is a classic *trans*- bond between Lys8 and Lys7, without interruption of the backbone H-bonds, displaying a continuous β-sheet between Asp4-Ile12 (Fig. 4A and S9). In addition, an interchain H-bond between Gly1 and Asp4 was formed (Fig. S9A), similarly to Pol III.

**Figure 4.**
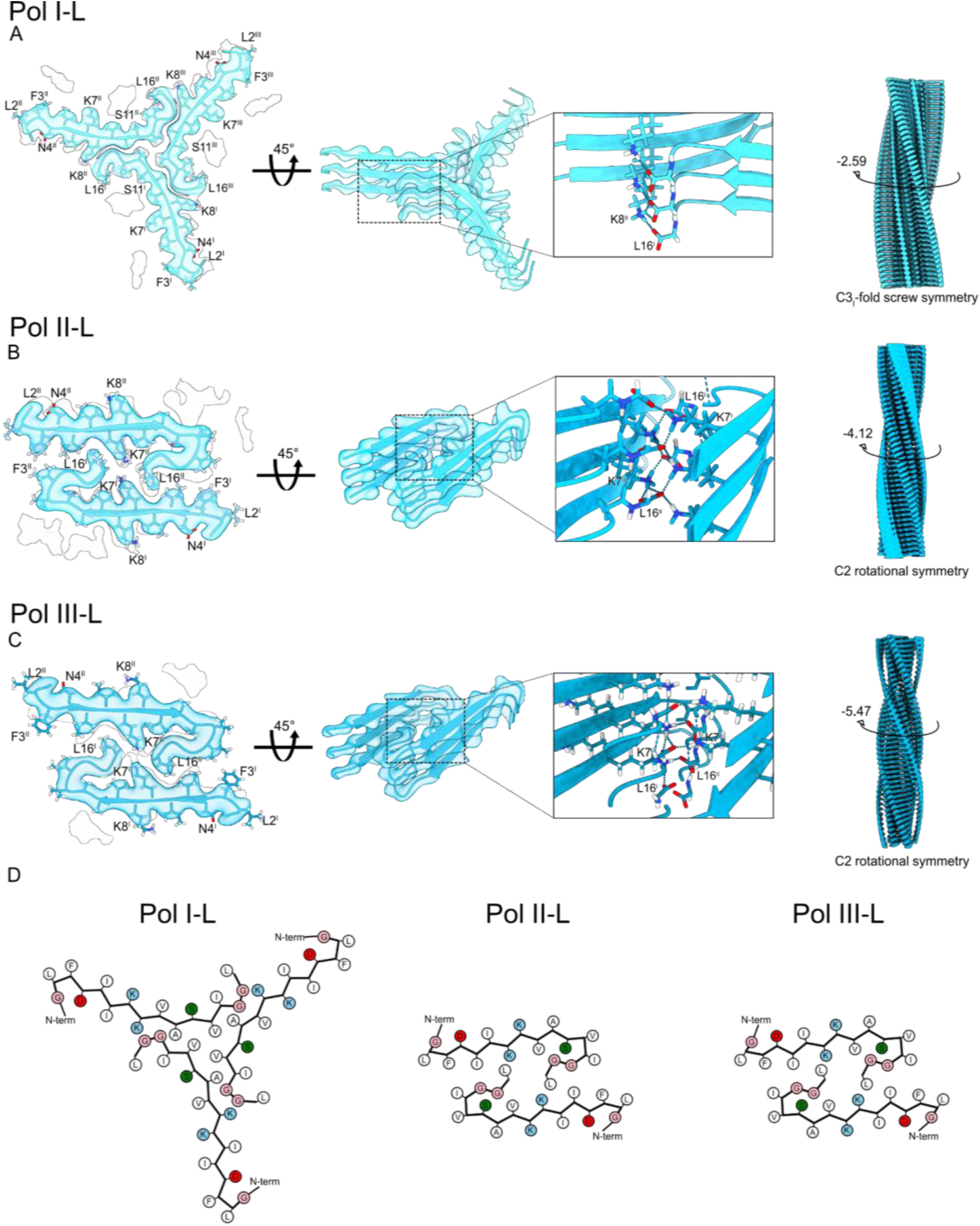
**Structure of citropin 1.3 at pH 7.4 in presence of liposomes**. **A)** Polymorph I-L displaying three peptide chain per layer, marked ^I,II,III^. **B)** Polymorph II-L and **C)** Polymorph III-L displaying two peptide chain per layer, marked ^I,II.^ The three polymorphs contain potential additional protein chains suggested by an incomplete map flanking the resolved fibril part. **A-C)** Left panels: atomic model of a single layer built into the Coulomb density with visualization of low-resolution densities. Middle panels: Three-layered density map featuring the fit of the atomic model, accompanied by close-up insets highlighting the formed hydrogen bond between Leu16 and Lys8^I^ (**A**) or Leu16 and Lys7^I^ (**B&C**). Left panels: full maps with reported twist. **D)** Cartoon of residue properties within the fibril cross-section. Hydrophobic, polar, and negatively/positively charged residues are indicated in white, green, red, and blue, respectively.

The second and third fibril polymorph of the dataset, Pol II-L and III-L, presented identical morphologies which differed only in twist and crossover (Table 1). Pol II-L and III-L had a resolution of 3.20 and 3.58 Å, respectively, with C2 symmetry from which two chains per layer could be modelled (Fig. 4B,C) with loose extra densities similar to Pol I-L (Fig. S8). For both maps, the fitted peptide presented an extended β-sheet between Asp4 and Ile10 with a LARKS kinked motif at residues Val12 and Leu13 where a *cis*-bond and angles typical of α-helices were formed (Fig. S10A, S11A). In the core of the structure, the amine groups of Lys7, and the C-terminal carboxyl groups of Leu16 formed crossed salt bridges resulting in a very tight interaction. Differently from other polymorphs, Pol II-L and lII-L showed a marked twist of -4.12° and -5.47°, respectively with a strong in-plane chain tilt (Fig. S9, S10). Overall, a larger fibril twist is observed in the polymorphs grown in the presence of liposomes.

**Table 1.**
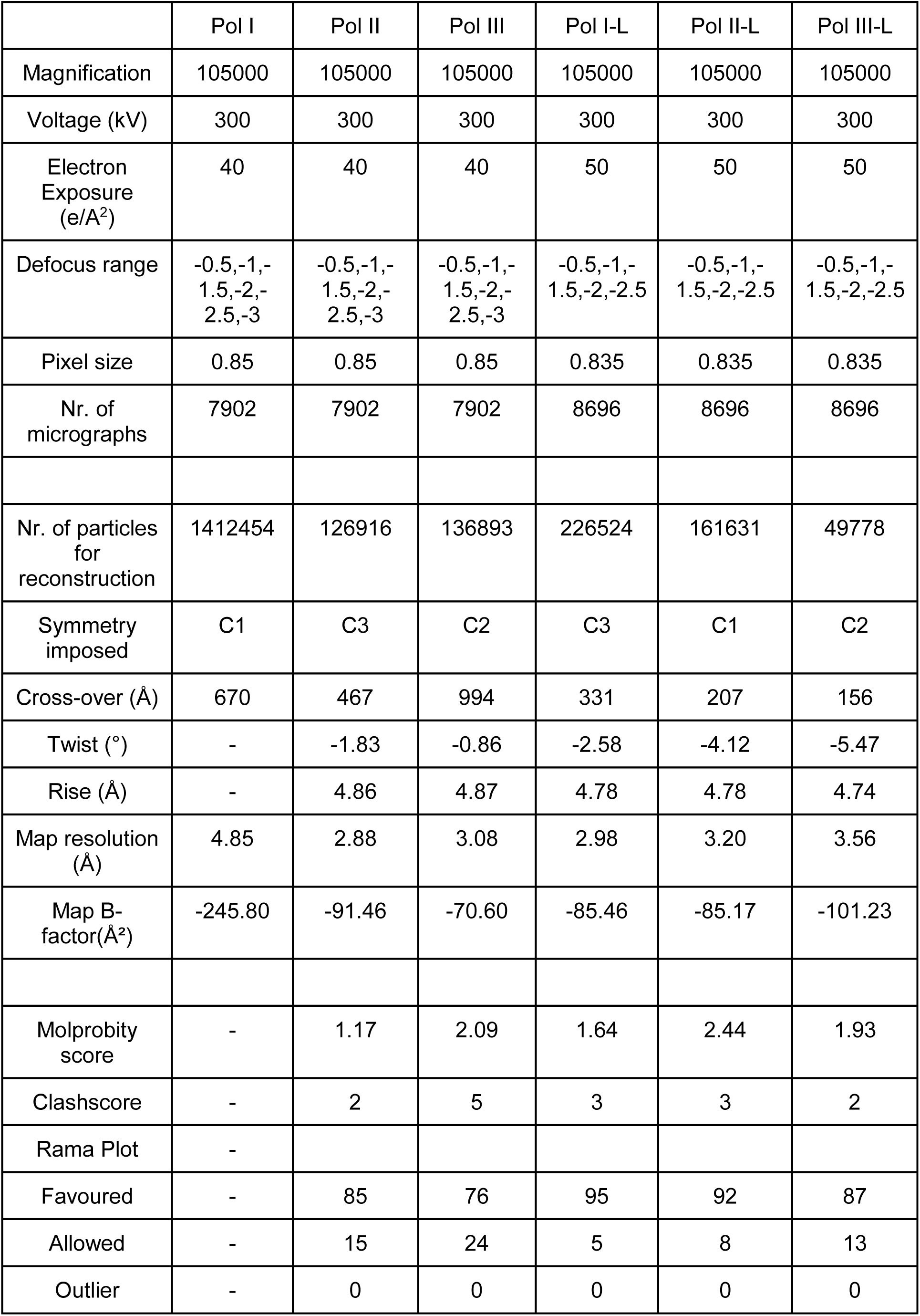
Cryo-EM Data collection parameters and map/structure statistics.

### Citropin 1.3 induces lipid vesicle fusion and undergoes liquid-liquid phase separation

The interaction of citropin 1.3 with membranes was explored using fluorescently labelled giant unilamellar vesicles (GUVs) with a net negative charge ^29,30^ (composition is detailed in the Method section), as a simple model system of a Gram-positive bacteria. The advantage of using GUVs is the possibility to follow a single membrane bilayer and therefore understanding the influence of citropin 1.3 on membrane integrity. Citropin 1.3 was labelled with fluorescein isothiocyanate (FITC) at the N-terminal end, which retained fibril forming ability (Figure S12), and the interactions were monitored via fluorescence microscopy in a time course experiment (Fig. 5).

**Figure 5.**
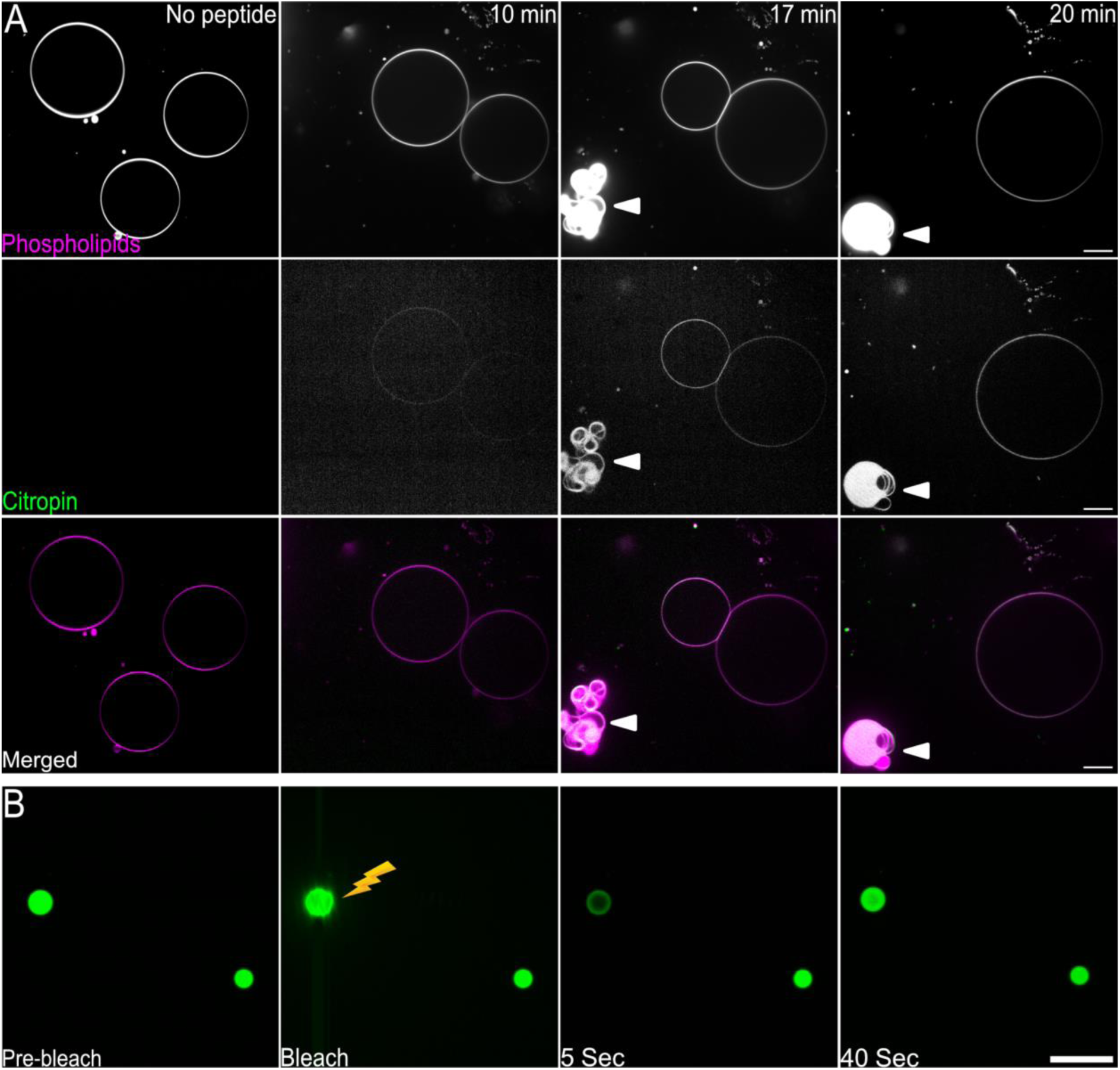
Interactions between negatively charged GUVs and citropin 1.3 monitored by fluorescence microscopy. **A)** Time dependent (0, 10, 17 and 20 min) experiments of the interaction between GUVs with a net negative charge (phospholipids panel, colored in magenta in the merged panel) and FITC-labelled citropin 1.3 (citropin panels, colored in green in the merged panel). The addition of the citropin 1.3 induced GUV fusion and the formation of condensates compose of the phospholipids and the peptide (indicated with white triangle). Scale bars, 10 μm. **B)** FRAP experiment targeting a single droplet (lightning symbol) displaying intensity recovery after bleaching, thereby supporting LLPS. Scale bar, 10 μm.

In absence of the peptide (t = 0 min), big spherical GUVs were visible (Fig. 5A). After the peptide injection and diffusion in the well, the peptide slowly started to accumulate on the membrane (t = 10 min), inducing the merging of neighboring GUVs into a single larger vesicle (t = 20 min). Such fusion based mechanism causes the permeabilization of membranes, a mechanism that differs from the classical lytic effect on membranes exploited by similar peptides ^20^. In the last two time points (t = 17 min and t = 20 min), it was possible to observe the formation of droplets as indicated by the triangular markers (Fig. 5A). This indicates the incorporation of the FITC-labelled peptide into condensates. The liquid properties of these condensates were subsequently confirmed by fluorescence recovery after photobleaching (FRAP) (Fig. 5B and S13), supporting liquid-liquid phase separation (LLPS) ^31^.

Citropin 1.3-mediated fusion into larger vesicles was also observed for liposomes composed of multilamellar and large vesicles prepared via the thin film hydration method ^32^ (Fig. S14). However, no droplets were formed, suggesting that the LLPS of GUVs and citropin 1.3 might have been influenced by the agarose used during GUV preparation, potentially acting as a crowding agent to facilitate phase separation.

### Citropin 1.3 exhibits toxicity to mammalian cells and induces LLPS with cellular components

The lethal concentration of citropin 1.3 required to kill 50% of human lung epithelial cells (A-549) (LC50) was determined to be 21 μM (Fig. S15). The same concentration was subsequently used for live-cell imaging to monitor the effects of citropin 1.3 on A-549 cells via fluorescence microscopy. A-549 cells were stained with wheat germ agglutinin to label cell membranes, Hoechst 33342 to stain DNA and the nucleus, and propidium iodide to identify dead cells and nucleic acids. Citropin 1.3 was applied as a mixture of 1% FITC-labeled and 99% unlabeled peptide.

Ten minutes after the addition of citropin 1.3 monomers, the peptides began accumulating on the plasma membrane, leading to cell death and membrane blebbing (Fig. 6A). Following membrane permeabilization and cell death, citropin 1.3 accumulated intracellularly, particularly within the nucleolus, a membraneless nuclear organelle involved in ribosome biogenesis ^33,34^. The colocalization of FITC and PI fluorescence intensities was analyzed by plotting their normalized pixel intensities against one another, yielding a Pearson’s correlation coefficient of 0.802 (Fig. S16). This high correlation confirms the colocalization of the two dyes ^35,36^, further supporting the accumulation of citropin 1.3 in the nucleolus. At later time points (t = 240 min), droplet-like structures were observed forming in the cytosol (Fig. 6B). FRAP experiments confirmed that citropin 1.3 induced LLPS with cellular components (Fig. S17).

**Figure 6.**
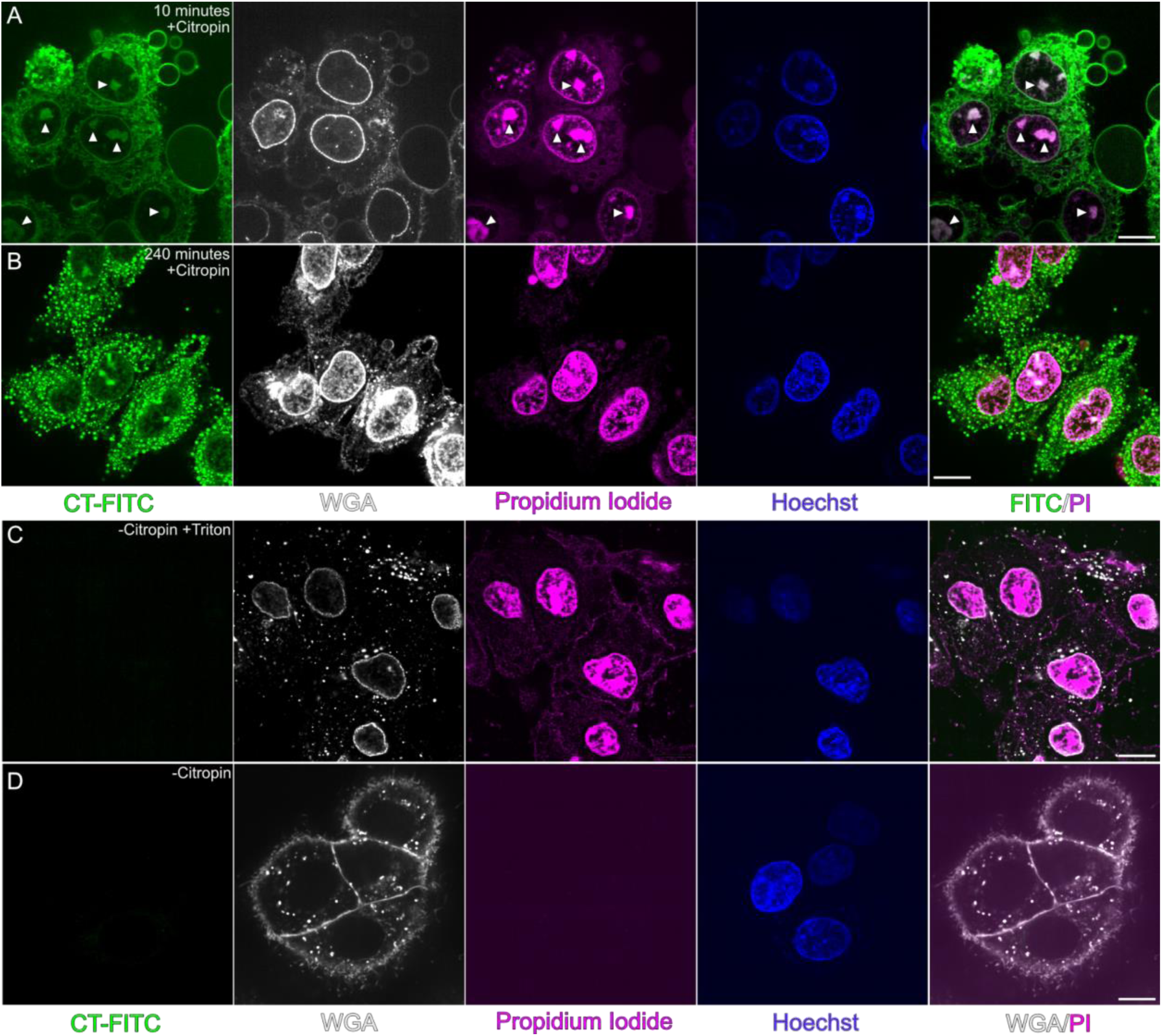
Citropin 1.3 acting against mammalian cells. Fluorescence live cell microscopy visualizing interactions between 21 μM citropin 1.3 and lung epithelial cells (A-549), tracked using four different fluorophores: citropin 1.3 (1% FITC-labelled and 99% unlabeled), wheat germ agglutinin (WGA), propidium iodide (PI) and Hoechst 33342 with colors reported in legend. **A)** Cell death induced by citropin 1.3 after 10 minutes of its injection as indicated by positive propidium iodide fluorescence. The colocalization of citropin 1.3 and PI-bound cellular components (i.e., containing nucleic acids) is indicated by white at the FITC and PI channels. **B)** A frame captured 240 minutes after the addition of citropin 1.3, showing the formation of peptide-rich droplets. **C)** Positive control showing cell death induced by Triton X. **D)** Negative control of cells without the addition of citropin 1.3. Scale bar: 10 μm.

The activity of citropin 1.3 against the Gram-positive bacterium *Bacillus subtilis* was quantified, with the minimum inhibitory concentration (MIC) determined to be 8 μM (Fig. S15). This concentration was subsequently used to visualize the live interaction of citropin 1.3 with bacterial cells using fluorescence microscopy. Initially, the peptide bound to the bacterial plasma membrane (Fig. S18), and through a membrane permeabilization mechanism, it induced bacterial cell death, as evidenced by the penetration of propidium iodide (Fig. 7A). Over time, citropin 1.3 accumulated in the bacterial cytosol, with only limited colocalization observed between the peptide and the bacterial genetic material (Fig. 7A).

**Figure 7.**
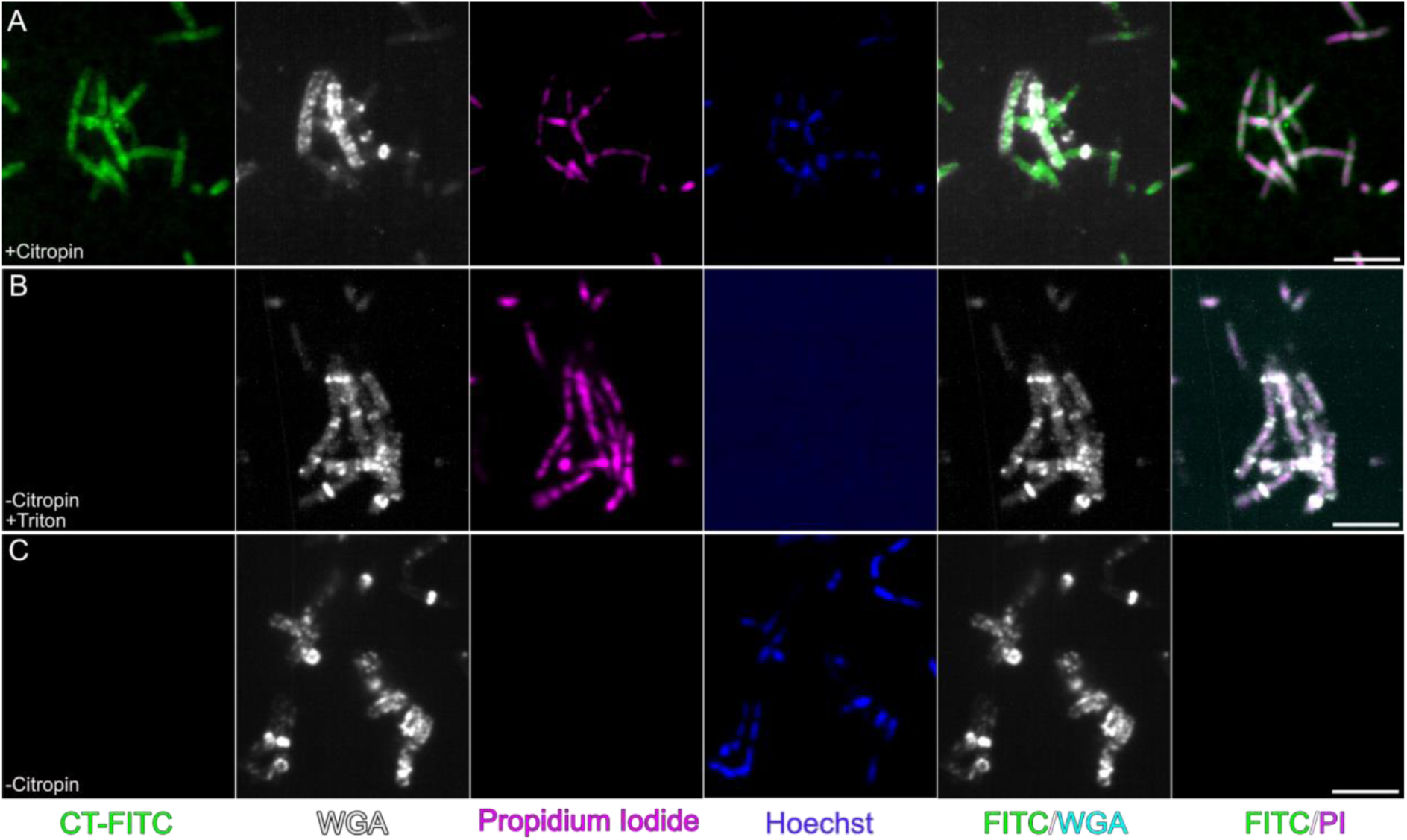
Citropin 1.3 acting against *Bacillus subtilis*. Fluorescence live cell microscopy visualizing interactions between citropin 1.3 at 8 μM and *Bacillus subtilis*. **A)** Bacterial death induced by citropin 1.3 is indicated by positive propidium iodide fluorescence with internalization in cell cytosol. **B)** Positive control with Triton X and consequent bacterial death, **C)** negative control of cells with no peptide added. Scale bar, 5 μM.

## Discussion

AMPs constitute a broad and extensively studied class of biomolecules, widely regarded as promising alternatives to conventional antibiotics in combating the global crisis of drug-resistant pathogens ^37^. The activity and regulation of AMPs are governed by diverse mechanisms, one of which involves a recently proposed hypothesis linking their function to amyloid fibril formation ^38^. This idea is supported by observations that certain pathological human amyloids, associated with neurodegenerative diseases, exhibit antimicrobial properties ^6,39,40^. Our findings on citropin 1.3 provide compelling evidence of a functional and structural connection between its antibacterial activity and the formation of amyloid fibrils. These results highlight the importance of further investigating this relationship, particularly in the context of the physiological roles of human amyloids. Additionally, a deeper understanding of the SAR in amyloidogenic AMPs is crucial for the rational design and development of more effective antimicrobial agents.

Cryo-EM analysis of citropin 1.3 revealed the presence of multiple fibril morphologies, each forming under distinct aqueous conditions. This phenomenon of amyloid polymorphism is well-documented across numerous amyloid structures, displaying a different energy landscape compared to globular proteins. Such polymorphism reflects kinetic trapping driven by environmental factors, interacting molecules, and inherent variability even within the same sample ^41–45^. Our findings identified at least ten distinct fibril morphologies, ranging from canonical cross-β amyloids to multi-layered nanotubes, and a crystal structure of a supramolecular α-helical structure, overall illustrating remarkable quaternary polymorphism.

Five high-resolution cryo-EM structures of citropin 1.3 fibrils were resolved, all exhibiting cross-β amyloid architecture with distinct lateral arrangements. Pol II-L and Pol III-L shared the same peptide fold, differing only in fibril crossover and twist. Such variations have also been observed in time-dependent studies of human islet amyloid polypeptide (hIAPP), suggesting that differences in crossover patterns may correspond to fibril transition states ^42^. The citropin 1.3 polymorphs demonstrated both fully extended and kinked β-sheets. This intrinsic chain flexibility is likely attributable to the presence of three glycines within the 16-residue peptide sequence. Notably, a kink in the β-strands of Pol II-L and Pol III-L occurred at Gly14-Gly15 (Fig. 4B,C). In contrast, Pol II exhibited a kink around Lys8-Val9 (Fig. 1), while Pol I-L displayed a kink near Val12-Ile13, preceding the double glycine motif (Fig. 4A). These structural variations emphasize the adaptability of citropin 1.3 fibrils and their potential for diverse functional and mechanical properties.

A key interaction observed across all resolved polymorphs was the conserved hydrogen bond formed between the positively charged amine of Lys7 or Lys8 and the negatively charged C-terminal carboxyl group of Leu16 on an adjacent chain. This electrostatic bond, consistently present in each fibril layer, appears to play a critical role in stabilizing the fibril architecture. Additionally, nearly all polymorphs exhibited an intra-chain hydrogen bond between Asp4 and Gly1, except from Pol II, in which the N-terminal end is extended (Fig. 1A). These findings underscore the importance of both inter- and intra-chain interactions in maintaining fibril stability and highlight the structural variability among citropin 1.3 polymorphs.

An abundant polymorph, observed at pH 5 but limited by resolution constraints preventing unambiguous modeling, is Pol I nanotube structure (Fig. 2). This polymorph displayed a quaternary arrangement of three concentric layers, speculated to include multiple secondary structure assemblies. Features such as a crossover distance of 670 Å (Fig. 2B), reflections at 4.75 Å in the power spectra (Fig. 2A), and densities in the map resembling β-sheets in the middle layer strongly suggest amyloid-like properties. Additionally, the reconstructed map’s outer layer (Fig. 2C) suggests the presence of α-helices wrapping and decorating the amyloid core. This unique fibril arrangement simultaneously incorporates α- and β-secondary structure elements, representing an intriguing dual-structural amyloid organization. Nanotube structures with two to four layers were also observed under neutral pH conditions (Fig. S2). Other amyloid-forming peptides, such as the phenol-soluble modulins PSMα3 and PSMβ2 secreted by *Staphylococcus aureus*, have previously been shown to form nanotube structures, as revealed by cryo-EM. These nanotubes consist of a cross-α arrangement of mated α-helical sheets that further assemble into tubular architectures ^46,47^. While the cross-α arrangement of PSMα3 was initially resolved using X-ray micro-crystallography ^6^, its supramolecular nanotube assembly was only discernible via cryo-EM ^44^. The shared tendency of citropin 1.3 and PSMs to form nanotubes hints at a potential connection in their SAR. However, the precise role of these nanotubes in cell membrane disruption remains unclear, warranting further investigation.

Our findings also demonstrated that the addition of negatively charged liposomes to citropin 1.3 monomers shifted the aggregation preference from nanotubes to cross-β amyloids (Fig. 4). This suggests that cell membranes may influence the peptide’s aggregation pathways and supramolecular arrangements. Fluorescence confocal microscopy studies using different membrane models revealed that citropin 1.3 induced the fusion of GUVs (Fig. 5). This mechanism differs from the classical membrane lysis typically associated with cationic AMPs ^48–50^. Membrane fusion mediated by citropin 1.3 might be a consequence of peptide insertion into the membrane, causing deformation and bringing opposing membranes into close proximity ^51,52^. This fusion mechanism may be linked to *in vivo* membrane permeabilization and subsequent cell death, offering a novel perspective on the antimicrobial activity of citropin 1.3.

Citropin 1.3, with a net positive charge of +1 at neutral pH, primarily consists of hydrophobic residues (9 out of 16 residues) and adopts an amphipathic α-helical structure in its monomeric form. It is hypothesized to interact with negatively charged bacterial membranes, leading to their disruption or permeabilization ^48–50^. Both the cross-β polymorphs and the crystal structure featuring α-helices revealed surfaces characterized by hydrophobic and mostly positively charged patches (Figs. 3 and S19). These patches may serve as sites for interaction with other peptides, as suggested by additional densities in the cryo-EM maps (Figs. 1 and 4), or potentially interact directly with membrane lipids. A similar structural pattern was observed in the active fragment of the human antimicrobial peptide LL-37, specifically residues 17–29 ^8^. Yet, this active fragment is more positively charged (+4 at neutral pH) than citropin 1.3, which might lead to differences in the mechanisms of membrane disruption.

Simultaneously with peptide-induced membrane fusion, we observed the formation of liquid condensates, which appeared exclusively in the presence of agarose, likely acting as a crowding agent (Figs. 5 and S13). The formation of liquid condensates has been associated with accelerated amyloid fibril aggregation, as these condensates create localized environments of high peptide concentration ^53–55^. Given that both bacterial and eukaryotic cells secrete macromolecules and polymers that can function as crowding agents, it is plausible that such condensate-driven effects may occur under physiological conditions. In fluorescence live-cell microscopy experiments, citropin 1.3 rapidly interacted with the membranes of A-549 mammalian cells, leading to cell death (Fig. 6). Intriguingly, after cell permeabilization, the peptide accumulated in the nucleolus, a membraneless organelle rich in proteins, DNA, and RNA ^56^. This localization is notable as RNA-amyloid interactions are known to facilitate LLPS, which can modulate protein aggregation ^57–59^. For example, nuclear proteins undergoing LLPS in response to stress or DNA damage can transition to fibril formation when dysregulated or mutated. Such processes are linked to neurodegenerative diseases like amyotrophic lateral sclerosis and frontotemporal dementia ^60–62^. The accumulation of citropin 1.3 in the RNA-rich nucleolus aligns with the observed formation of peptide condensates (Fig. 6A, t = 240 min). AMPs have previously been proposed to combat bacterial infections by targeting nucleic acids through phase separation, compacting them to inhibit transcription and translation ^63^. Our findings connect this mechanism to amyloid fibril formation, whether on- or off-pathway to LLPS. These results open new perspectives on the potential of small amphiphilic peptides like citropin 1.3 to form heterotypic fibrils with genetic material. This capability could have implications for immunomodulation and developing novel strategies for combating infections, particularly by targeting RNA or DNA-rich compartments.

Fluorescence microscopy of Bacillus subtilis treated with citropin 1.3 demonstrated that the peptide effectively killed bacterial cells by disrupting the plasma membrane (Fig. 7). Notably, no overlap was observed between FITC-labeled citropin 1.3 and the WGA dye (Fig. 7A), which specifically binds to cell wall components ^64^. While some intracellular accumulation of the peptide was detected, minimal and sporadic colocalization with bacterial genetic material was observed, and no evidence of LLPS was detected. The observed differences in peptide localization and behavior between eukaryotic and prokaryotic cells suggest distinct mechanisms of cytotoxicity and interactions with cellular components. In eukaryotic cells, the nucleolus appears to provide a favorable environment for interactions with citropin 1.3, potentially facilitating nucleic acid-induced LLPS. In contrast, in bacterial cells, the absence of such interactions indicates that citropin 1.3 primarily exerts its antimicrobial effects via membrane disruption rather than through interactions with intracellular components. These findings underscore the context-dependent activity of citropin 1.3 and highlight its distinct modes of action in different cell types, emphasizing the peptide’s versatility as both an antimicrobial agent and a mediator of cellular phase separation under specific conditions.

## Conclusions

Overall, our findings reveal the polymorphic nature of citropin 1.3 amyloid structures, including cross-β fibrils and multi-layered nanotubes likely composed of distinct secondary structure elements, displaying unique structural assemblies. Our results further suggest that membranes play a significant role in promoting fibrillation into the cross-β amyloid form. This interaction mechanism appears distinct from that of uperin 3.5, where lipids stabilize an α-helical conformation^6^. While citropin 1.3 fibrils themselves may not directly mediate cytotoxicity, the fibrillation process likely contributes to regulating the peptide’s activity and target-cell specificity. Additionally, our findings indicate that certain macromolecules in eukaryotic cells can trigger LLPS of citropin 1.3. LLPS was also observed in the presence of phospholipids and a crowding agent, highlighting the role of environmental factors in modulating peptide behavior. Furthermore, the observed colocalization of citropin 1.3 with genetic material and the formation of condensates in eukaryotic cells suggest a functional interaction between the peptide and nucleic acids. This interaction may play a role in regulating the peptide’s activity, linking its structural versatility to diverse biological functions. Together, these insights highlight the complex interplay between citropin 1.3’s structural polymorphism, its environmental context, and its biological activity, offering a deeper understanding of its multifunctionality.

## Methods

### Peptide aggregation

Citropin 1.3 from *Litoria citropa* (Southern bell frog) (Uniprot ID P81846; sequence GLFDIIKKVASVIGGL) and citropin 1.3 labeled with FITC at the N-terminal end were purchased from GL Biochem Ltd. (Shanghai) as lyophilized peptides, at >98% purity. The peptide was incubated in double distilled water (ddH2O) in 1.5 mL tubes at room temperature without shaking. The optimal incubation time to produce ordered and dispersed fibrils was monitored over time by a negative-stain transmission electron microscopy. This time point was used for the preparation of cryo-EM samples.

### Liposome and GUVs preparation

Two distinct methods were used for the preparation of membrane model systems: Liposomes of multilamellar and large unilamellar vesicles, composed of 2:1 molar ratio of 1,2-dioleoyl-sn-glycero-3-phosphoethanolamine (DOPE): 1,2-dioleoyl-sn-glycero-3-phosphoglycerol (DOPG), were prepared using the thin layer method ^65^. For the large unilamellar vesicles formation, an extruder device (Avanti) with 500 nm filters was used on multilamellar vesicles. These liposomes were used for the cryo-EM studies in Figure 4 and for the fluorescence microscopy experiments in Figure S14. For the fluorescence microscopy experiments, 1% of total lipid mass of 18:1 Liss Rhod PE (Avanti) was added to each mixture. GUVs, used for the fluorescence microscopy experiments in Figure 5, were composed of 2:1 (molar ratio) DOPE:DOPG, prepared as in the agarose swelling method ^66^.

### Cryo-electron microscopy

For cryo-electron microscopy, citropin 1.3 was incubated in three different conditions: I) 1 mM citropin 1.3 in 50 mM NaCl at pH 5 and 48 h incubation; II) 1 mM citropin 1.3 in 10 mM phosphate-buffered saline (PBS) at pH 7.4 and 48 h incubation; III) 500 μM citropin 1.3 in 10 mM PBS at pH 7.4 in the presence of 2.5mM negatively charged 1:1 DOPE:DOPG liposomes and 24 h incubation.

For each condition, 3.5 μl of sample were deposited on glow-discharged holey carbon grids (Quantifoil R2/1, 300 mesh) at 100% humidity and 20 °C, blotted after 10 s of wait time with empirically optimized parameters, and subsequently plunge-frozen in liquid ethane-propane using a FEI Vitrobot Mark IV (FEI microsystems). Samples were imaged on a Titan Krios G3i (ThermoFisher Scientific) transmission electron microscope, operated at 300 kV and equipped with a Gatan Bioquantumenergy filter operated in zero-loss mode (20 eV energy slit width). Images were acquired on a Gatan K3 electron counting direct detection camera (Gatan Inc.) in dose fractionation mode using EPU software at a nominal magnification of 105000x. Data collection details are described in Table 1.

Fibrils of 0.5 mM FITC-labeled citropin 1.3 incubated for 24 hours in water were imaged using a ThermoFisher Scientific (FEI) Talos F200C transmission electron microscope, operating at 200 kV and equipped with a Ceta 16M CMOS camera, at the Ilse Katz Institute for Nanoscale Science and Technology, Ben Gurion University of the Negev, Israel.

### Image processing and polymorph separation

The initial motion correction and CTF estimation of the motion-corrected micrographs was performed using the MotionCorr2 ^67^ and CTFFIND 4 ^68^ implementation in Relion 5 ^69,70^. Automatic segment picking with inter-segment distance of 300 pixels and filament tracing was then performed in crYOLO ^71^, and found segment coordinates were exported in *helix* format. Using RELION 5, 2D class averages were computed as a first step of polymorph identification. For each identified polymorph the built-in *relion_helix_inimodel2d* was used for generating initial models for following 3D refinements using Blush regularisation in Relion ^70^ . The final volumes were obtained from RELION 5 for all polymorphs except for the polymorph I whose final volume was obtained using cryoSPARC 4 ^28^. From the obtained densities, the final volume used for model building was obtained using automatic B-factor sharpening in Relion.

### Model building

To build atomic models into the obtained maps, we proceeded identically for each polymorph: we created a single layered map, built a peptide monomer manually using Coot 0.9.377 ^72^ and then fitted inside a single helix layer. We then expanded this set of monomers into three full helix layers in ChimeraX ^73^ 1.8, using the symmetry parameters determined in map refinement. Interactive refinement guided by on-line molecular dynamics computations was then performed in ISOLDE 1.0b379 ^74^ within ChimeraX. Next, to mitigate edge effects of the molecular dynamics calculation, all helix layers but the central one were removed, and the single layer re expanded to three layers, as above. Model validation was carried out using MolProbity ^75^ within Phenix 1.19.180,81 ^76^. The obtained final model was further expanded to helices of arbitrary length using ChimeraX 1.8 for visualization, secondary structure analysis, Coulomb and lipophilicity potentials, with default parameters. Map-model per residue correlation coefficients show satisfactory values outside the aforementioned peripheral regions and flexible residues.

### Peptide crystallization

Citropin 1.3 was dissolved at 20 mg/ml in 20% acetic acid and ddH2O, mixed with a reservoir containing 20%v/v Glycerol, 8%w/v PEG 8,000 and 8%w/v PEG 1,000. Crystals were grown at 20 °C, using the hanging-drop vapor diffusion technique. 5% v/v 2-methyl-2,4-pentanediol (MPD) was added as cryo-protectant and single crystals were flash-frozen in liquid nitrogen before X-ray data collection. Data was collected at the EMBL P14 microfocus beamline at PETRA III at the Deutsches Elektronen-Synchrotron (DESY) on August 26, 2021.

### X-ray data processing, structure solution and refinement

Diffraction data from a single crystal was integrated and scaled in XDS to a resolution of 1.56 Å in space group P 4 3 2 with unit cell parameters (a=b=c= 67.83 Å). Dataset statistics can be found in Table 2.

**Table 2.** X-ray data collection parameters and structure statistics.

|  |  |
| --- | --- |
|  | Citropin 1.3 (pdb 9hpp) |
| **Data collection** |  |
| Space group | P 4<sub>1</sub> 3 2 |
| Cell dimensions |  |
| a,b,c (Å) | 67.83 , 67.83 , 67.83 |
| $\alpha\beta\gamma$ (°) | 90, 90 ,90 |
| Resolution (Å) | 47.97- 1.56 (1.66-1.56) |
| Rmeas (%) | 7.4 (240.7) |
| I/$\sigma$(I) | 14.65 (0.95) |
| Completeness (%) | 98.1 (90.5) |
| Redundancy | 9.56 (6.72) |
| Wilson B factor (Å<sup>2</sup>) | 29.08 |
| **Refinement** |  |
| Resolution (Å) | 47.96-1.56 (1.66-1.56) |
| Completeness (%) | 97.76 |
| No. reflections | 7856 |
| $R_{\text{work}}$ (%) | 28.39 (37.61) |
| $R_{\text{free}}$ (%) | 32.33 (38.31) |
| No. non-H atoms | 248 |
| Protein | 239 |
| Ligand/ion | - |
| Water | 9 |
| **B-factors** (Å<sup>2</sup>) |  |
| Protein | 34.19 |
| Ligand/ion | - |
| Water | 40.20 |
| **R.m.s. deviations** |  |
| Bond lengths (Å) | 0.014 |
| Bond angles (°) | 1.153 |
| All-atom clash score | 3.86 |
| Number of xtals used for scaling | One crystal |

Values in parentheses are for the highest-resolution shell.

Initial phases were recovered by molecular replacement with Phaser using the Phenix GUI. The NMR ensemble of aurein 1.2 (pdb 1vm5) supplemented with two copies of aurein 3.3 (determined in house, to be published) was used as the basis of the search model. The 11 N-terminal residues were kept, the terminal residue was trimmed down to its Cγ atom and the last isoleucine was trimmed down to a valine to better match the citropin 1.3 sequence. The ensemble consisting of 7 models was finally realigned prior to being used as a search model. A single copy was found in space group P 41 3 2 and the resulting map allowed for iterative manual model building and refinement in Coot and Phenix.refine respectively. The maps revealed a second copy of the citropin 1.3 peptide that could be fully built. Towards the center of the unit cell, exposed to the hydrophobic side of the clearly defined citropin 1.3 peptides is residual density possibly from a low occupancy or disordered third copy which could not be unambiguously built.

At the position where this third copy would be, it would not contribute to the propagation of the observed crystallographic lattice. This helps explain the poor interpretability of the observed electron density and resulting sub-optimal match between the observed and model derived structure factors as represented in the Rwork and Rfree values.

Reprocessing of the diffraction images while lowering the Bravais lattice type from primitive cubic to either rhombohedral, primitive tetragonal or primitive orthogonal did not result in any improvement or segregation of the electron density of the third copy. One caveat of this approach was the low completeness of all lower symmetry datasets. Model and refinement statistics can be found in Table 2.

#### Minimum Inhibitory Concentration assay

Lyophilized citropin 1.3 peptide was dissolved in ddH2O water at 1 mM. The peptide stock solution was then diluted into the bacterial media (Mueller Hinton Broth media) in a sterile 96-well plate in order to have seven 2-fold dilutions of citropin. 10X PrestoBlue solution was added to the well to reach a 1X final concentration. *Bacillus subtilis* (Leibniz Institute DSMZ-German Collection of Microorganisms and Cell Cultures GmbH) bacterial cells were diluted to an optical density measured at 600 nm of 0.1. The diluted cell suspension was then dispensed into the well of the 96-well plate by diluting it 200-fold. The plate was then sealed, and the cell growth was followed every 5 minutes for 24 hours using a FLUOstar CLARIOstar (BMG LABTECH) plate reader by measuring PrestoBlue fluorescence (excitation: 545 nm; emission: 600 nm). Before each measurement, the plate was shaken for 10 seconds at 100 rpm using a double orbital shaking mode. For each condition, blank (media with PrestoBlue) was subtracted and the averaged PrestoBlue intensity was plotted as a function of time. The MIC was defined as the minimal concentration inhibiting bacterial growth. Experiments included control serial dilutions of peptide buffer alone and peptide alone without bacterial cells. Experiments were performed at least three times and each experiment contained at least two replicates for each condition.

#### Mammalian cell culture and cell toxicity

A-549 human lung epithelial cells (A-549) (ACC 107, DSMZ) were grown in RPMI 1640 medium containing L-glutamine, supplemented with 10% heat inactivated fetal calf serum, at 37⁰C in 5% CO2. For treatment with the peptide, cells at a concentration of 0.125M/mL at a 100 μL volume were incubated in a 96-well plate for 24 hours. On the day of the experiment, cells were washed twice with PBS and exchanged for 50 μL of fresh media. Citropin 1.3 was dissolved in ultrapure water, and the concentration was determined using a Nanodrop (ThermoFisher). The peptide stock was then diluted in RPMI media to the 2x of the starting desired experimental concentrations. A concentration range of 1-128 μM of peptide was tested, using plate serial dilution with a total final volume per well equal 100 μL. 50 μL of 2x concentrated peptide was added to each well already containing 50 μL of media, totaling 100 mL per well. The plate was incubated for 2 hours at 37⁰C in 5% CO2. After the incubation period, the plate was spun at 200 g for 10 minutes. Cell lysis was quantified using the lactate dehydrogenase release assay, according to the manufacturer’s instructions (LDH; Cytotoxicity Detection Kit Plus, Roche Applied Sciences). The absorbance was measured at 490 nm and the reference wavelength was 690 nm. The luminescent response was observed using a plate reader (Clariostar, BGM). The one-way ANOVA multiple comparison test was employed for the analysis of statistical significance between distinct experimental groups. Experiments were conducted independently three separate times, each with three technical replicates. GraphPad Prism 10 (GraphPad Software) was applied for all statistical analyses of the data.

### Fluorescence light microscopy analysis

Human (A-549) and bacteria (*Bacillus subtilis*) cells have been stained with 1 µg/mL wheat germ agglutinin (Thermo Fisher Scientific), and 1 µg/mL Hoechst 33342 (Thermo Fisher Scientific) in Hank’s buffered saline solution (HBSS) for 20 minutes at 37°C, 5% CO2 for membrane and nucleus staining, respectively. The samples were then washed twice with HBSS. Cells were kept in HBSS for imaging. 0.5 µg/mL of propidium iodide was added in the well before addition of citropin 1.3 (1% FITC-labeled citropin 1.3 and 99% unlabeled). *Bacillus subtilis* were fixed on the bottom of the well by coating the well with a 0.01% solution of Poly-L-Lysine (Merck). Images were recorded using a Nikon Ti2 equipped with Yokogawa CSU W1 spinning disc unit with two Hamamatsu sCMOS cameras and a 100×/1.45 NA oil immersion objective. Live cell imaging was performed in 8-well live cell imaging chambers. Experiments with eukaryotic cells have been performed in 5% CO2 environments at 37 degrees. Image processing was performed using ImageJ ^77^. Colocalization data analysis was performed by hand-drawing a “region of interest” (ROI) over an area of interest to then calculate signal intensity to be then used for the calculation of the Pearson’s correlation coefficient ^78^ which measures the pixel-by-pixel covariance in the signal levels. Values were then normalized and plotted accordingly. The data analysis was performed using the software Origin 2022b.

### FRAP experiment

Fluorescence recovery after photobleaching experiments were performed using a Nikon Ti2 equipped with a Yokogawa CSU W1 spinning disc unit with an Andor iXon EMCCD camera. In addition, the microscope was equipped with a FRAP unit from Rapp Optoelectronics. Image processing was performed using ImageJ^77^ and the data analysis was performed using the software Origin 2022b by normalizing each data point of each measurement.

## Supporting information

The Supplemental Information contains: Figures S1-S19

## Acknowledgements

M.L. acknowledges research support from the Israel Science Foundation, Grant No. 2111/20 the Cure Alzheimer’s Fund, the Forschungskooperation Niedersachsen – Israel, Volkswagenstiftung, No: 76251-4659/2022 (ZN 4042).

Funded/Co-funded by the European Union (ERC, FuncAmyloid, 101087140). Views and opinions expressed are however those of the author(s) only and do not necessarily reflect those of the European Union or the European Research Council. Neither the European Union nor the granting authority can be held responsible for them.

Y.B. is an ARISE fellow with funding from the European Union’s Horizon 2020 research and innovation programme under the Marie Skłodowska-Curie grant agreement No 945405.

We would like to thank Edward Egelman and Ravi R. Sonani for stimulating discussions over the nanotube structures observed by cryo-EM. We would like to acknowledge the staff of the ID23 and ID30A beamlines at ESRF (Grenoble, France) as well as the staff at the P14 beamline operated by the EMBL at the PETRA-III storage ring (Hamburg, Germany) for the beamtime allocation and their support. We would like to thank G. Bourenkov and G. Chojnowski for stimulating discussions regarding data processing, and Kay Diederichs for assistance and advice in data processing. We would like to thank Peleg Ragonis-Bachar and Eilon Barnea for help with crystallization and data collection, and the TCSB - Technion Center for Structural Biology for crystallization setup and visualization. We would like to acknowledge Alexander Upcher and the Ilse Katz Institute for Nanoscale Science and Technology, Ben Gurion University of the Negev, Beer-Sheva, Israel, for TEM analyses.

We acknowledge the use of OpenAI’s GPT-4 model to improve the writing quality of parts of this manuscript.

## Author Contributions

F.S. and M.L. conceived the experiments. F.S prepared the samples, collected and processed cryo-EM data. F.S. performed model building. F.S. imaged and processed fluorescence microscopy data. S.M. assisted with cryo-EM data processing. M.P.C. and A.G. conducted mammalian cell work. J.M. conducted MIC assays. M.L and E.G. supervised experiments. All results were discussed with all authors. F.S. wrote the manuscript with contribution from all the authors.

## Competing interests

The authors declare no competing interests.

## Notes

### Competing Interest Statement

The authors have declared no competing interest.

