## Supplementary material for "Structural and Functional Versatility of the Amyloidogenic Antimicrobial Peptide Citropin 1.3": The Supplemental Information contains: Figures S1-S19

Figures S1-S19

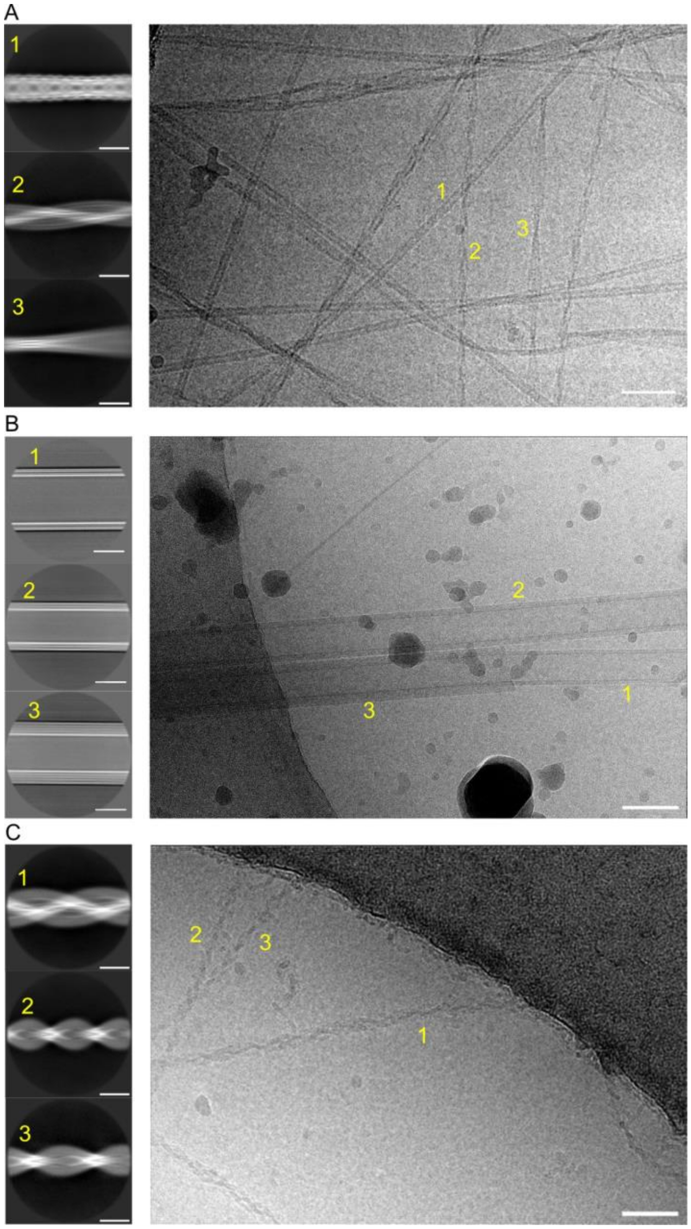


**Figure S1. Cryo-EM representative micrographs and 2D class averages of citropin 1.3 datasets from different incubation conditions**. **A)** 1 mM citropin 1.3 in 50 mM NaCl aqueous solution at pH 5, **B)** 1 mM citropin 1.3 in 10 mM PBS at pH 7.4 and **C)** 0.5 mM citropin 1.3 in 10 mM PBS in presence of 1.5 mM of DOPE:DOPG (2:1 molar ratio) liposomes. The different classes are numbered based on their relative particle count in each dataset. Scale bars, 500 Å for micrographs, 100 Å for 2D class averages in A) and C), 200 Å for 2D class averages in B).

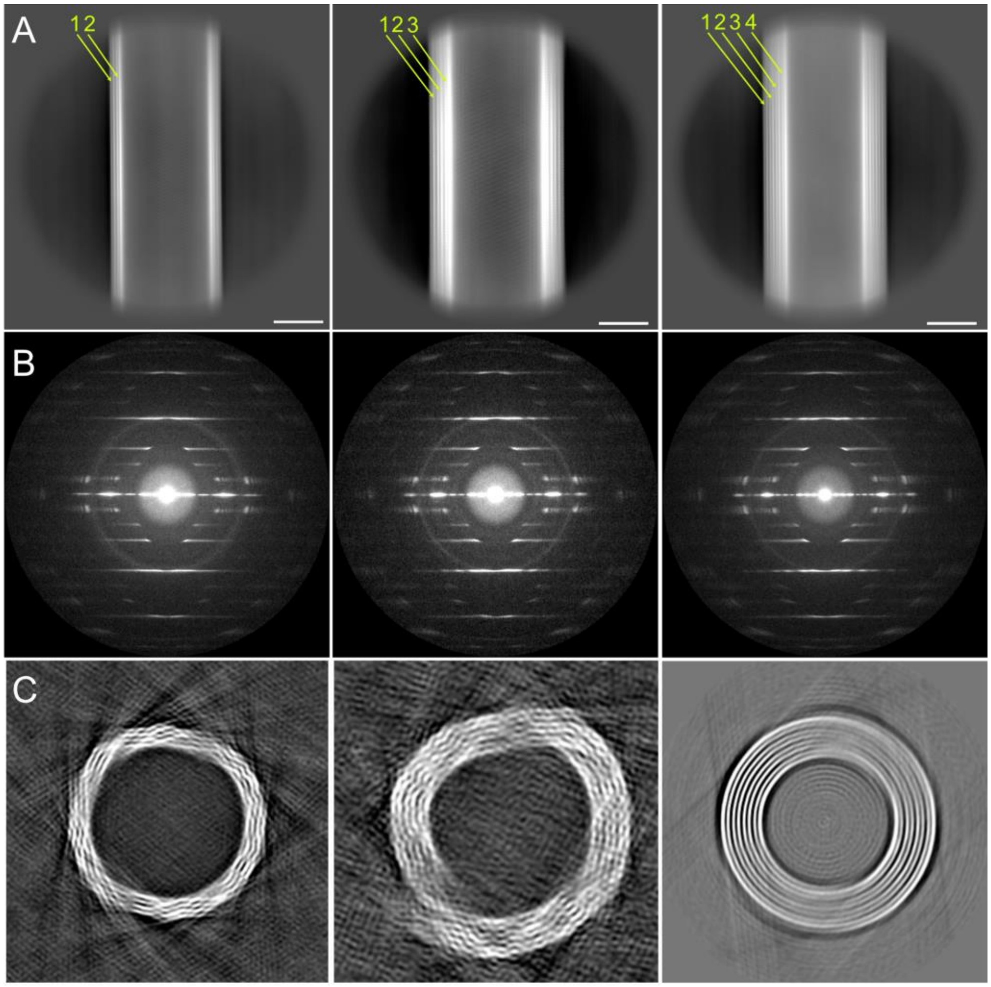


**Figure S2. Visualization of citropin 1.3 nanotubes formed in PBS at pH 7.4. A)** Representative cryo-EM 2D classes of obtained nanotubes with indication of number of layers for each class, scale bars are 200 Å. **B)** Power spectra corresponding to each 2D class. **C)** Cross-section of low-resolution 3D volumes obtained for each nanotube class.

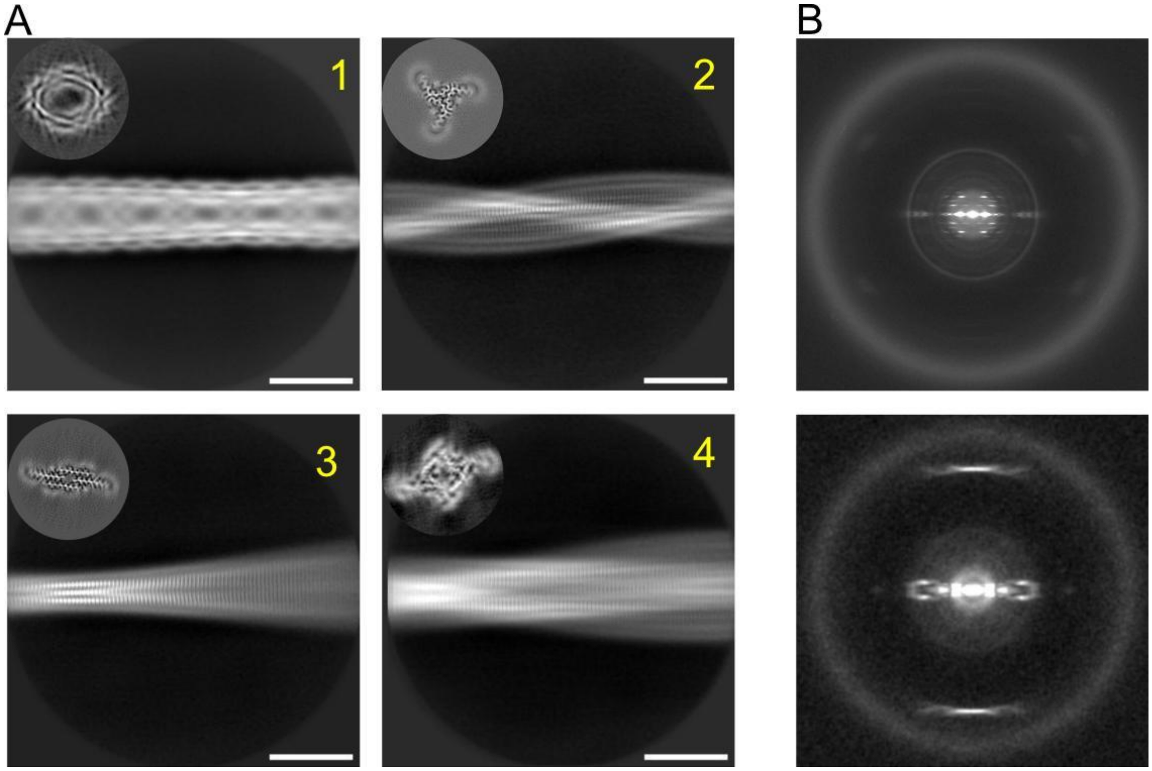


**Figure S3. Two dimensional details of citropin 1.3 fibrils formed at pH 5. A)** Cryo-EM 2D class averages of different polymorphs formed in 50 mM NaCl at pH 5, numbered 1 to 4, with 50.3%,16.2% 13.9%, 7.6% relative particle count per class on total percentage, respectively. The cross-sections of the relative class maps are shown as insets at the top right. Scale bars are of 50 Å. **B)** Obtained power spectra from Pol I (top) and representative amyloid class from Pol III (bottom).

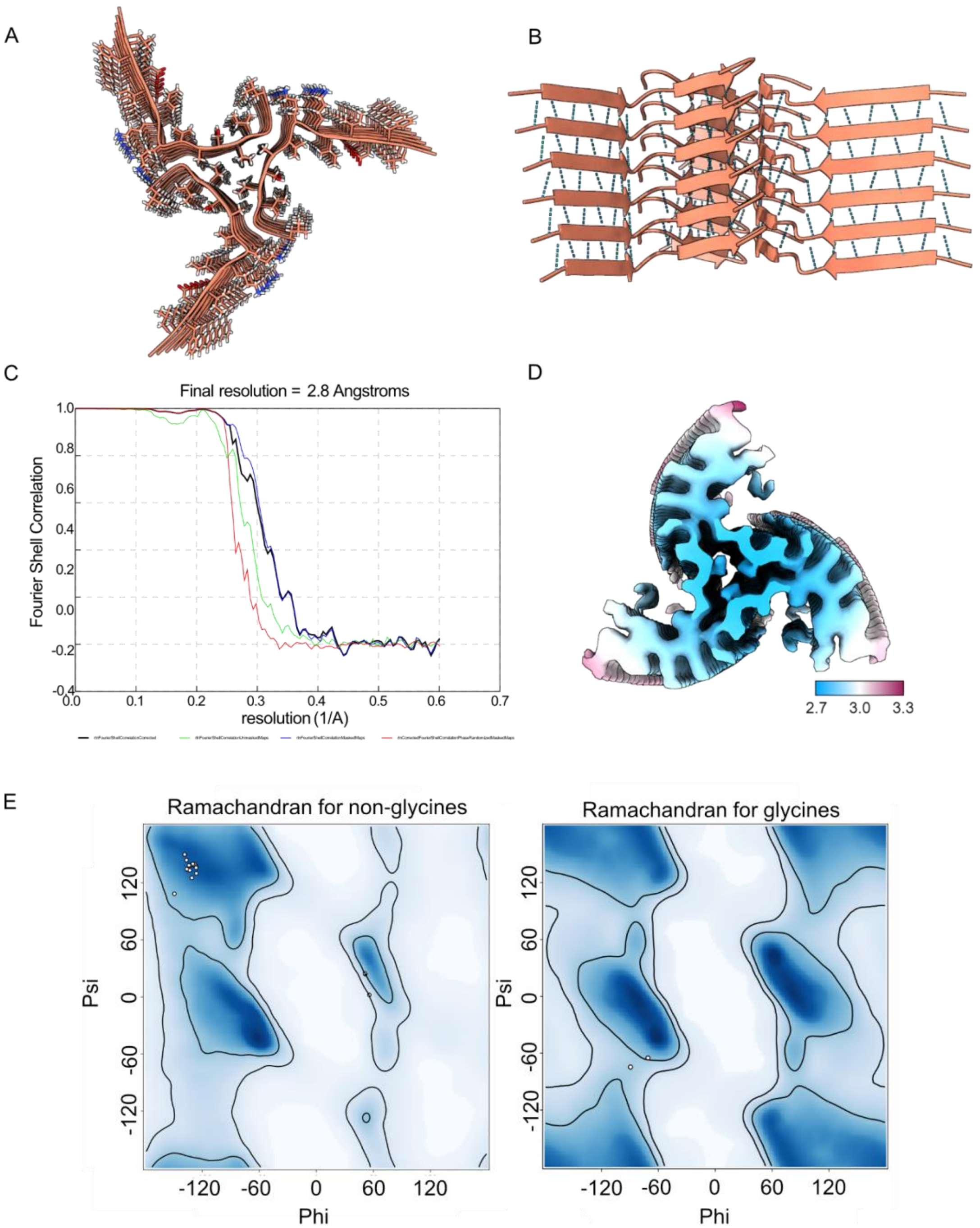


**Figure S4**. **Citropin 1.3 polymorph II cryo-EM map and model overviews. A)** A top view of the Pol II model with side chains shown as sticks. **B)** Side view with visualization of backbone hydrogen bonds. **C)** Gold Standard Fourier Shell Correlation (GSFC) curve reported from Relion of Pol II-L map and **D)** local resolution visualization of map cross-section. **E)** Ramachandran plots as reported from software Phenix for non-glycine and glycine residues. Most residues reside in the β-sheet region, except from Lys7 and Lys8, which display angles close to a left-handed α-helix.

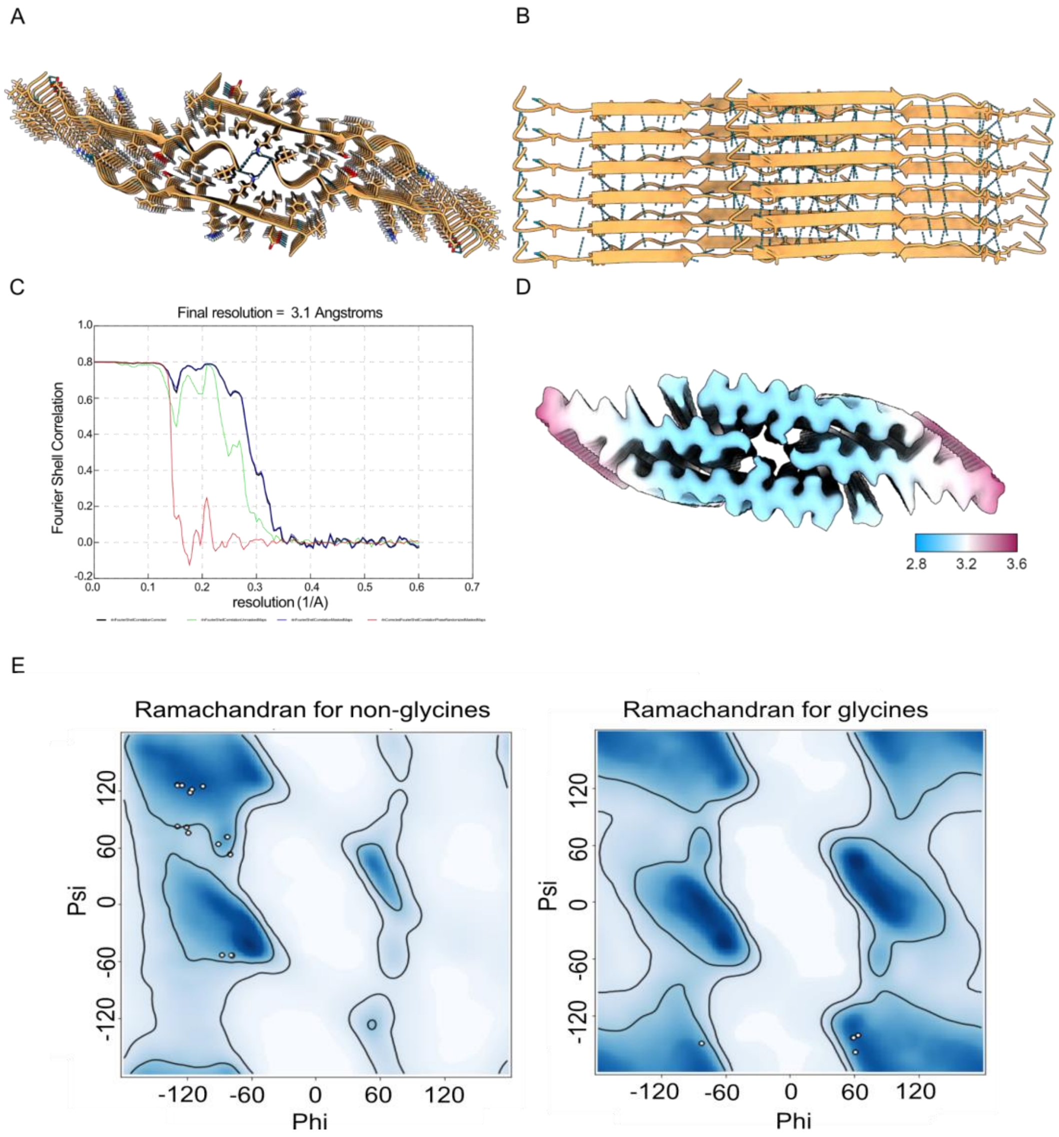


**Figure S5**. **Citropin 1.3 polymorph III cryo-EM map and model overviews. A)** A top view of the Pol III model with side chains shown as sticks. **B)** Side view with visualization of backbone hydrogen bonds. **C)** GSFC curve reported from Relion of Pol III-L map and **D)** local resolution visualization of map cross-section. **E)** Ramachandran plots as reported from software Phenix for non-glycine and glycine residues.

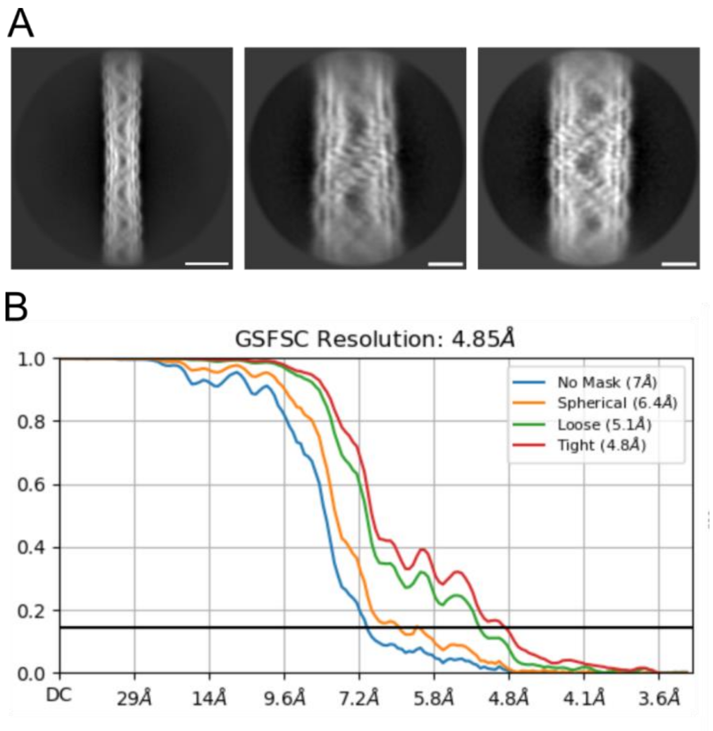


**Figure S6. Citropin 1.3 polymorph I 2D classes and 3D refinements. A)** Cryo-EM 2D classes of Pol I with 680 (left, scale bar of 100 Å) and 300 (middle and right, scale bar of 50 Å) pixels box size. **B)** Gold Standard Fourier Shell Correlation (GSFSC) curves with resolution reported by the CryoSPARC software.

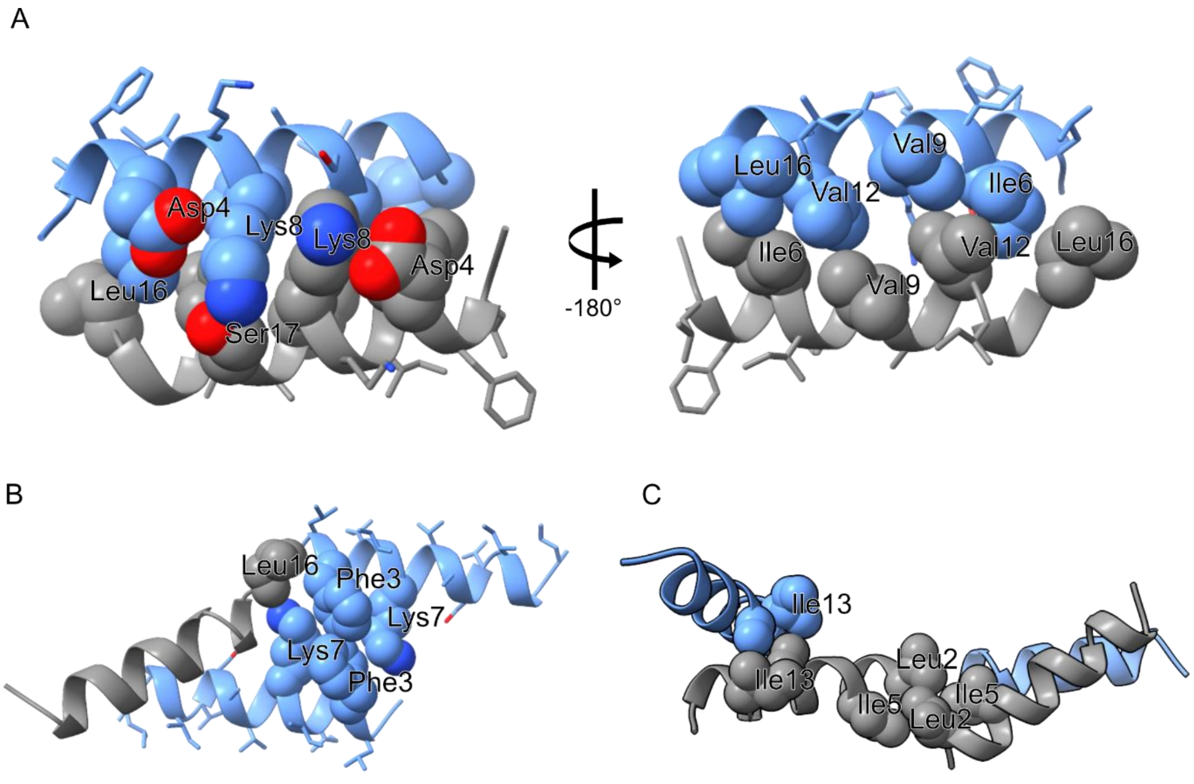


**Figure S7. Inter-α-helical packings in the crystal structure of citropin 1.3.** This figure highlights a subsection of the α-helical pairs in the citropin 1.3 crystal structure. Chains A and B are depicted in grey and light blue, respectively. **A)** The panel presents the interactions between the α-helices of chains A and B, shown in two orientations rotated approximately 180°. The helices exhibit tight packing mediated by hydrophobic interactions involving Ile5, Val9, Val12, and Leu16. Additionally, a hydrogen bond is observed between Lys8 on chain B and Ser11 on chain A. Each chain also features an internal electrostatic interaction between Lys8 and Asp4. **B)** The panel displays further stabilizing interactions between adjacent pairs, including tight packing between Phe3 and Lys7 of neighboring chain Bs, as well as interactions with Leu16 from chain A. **C)** The panel displays further stabilizing interactions between adjacent pairs, including tight packing between Leu2 and Ile5 of neighboring chain As, and between Ala10 and Ile13 of neighboring chain A and B. These interactions collectively contribute to the stability and organization of the α-helical assembly.

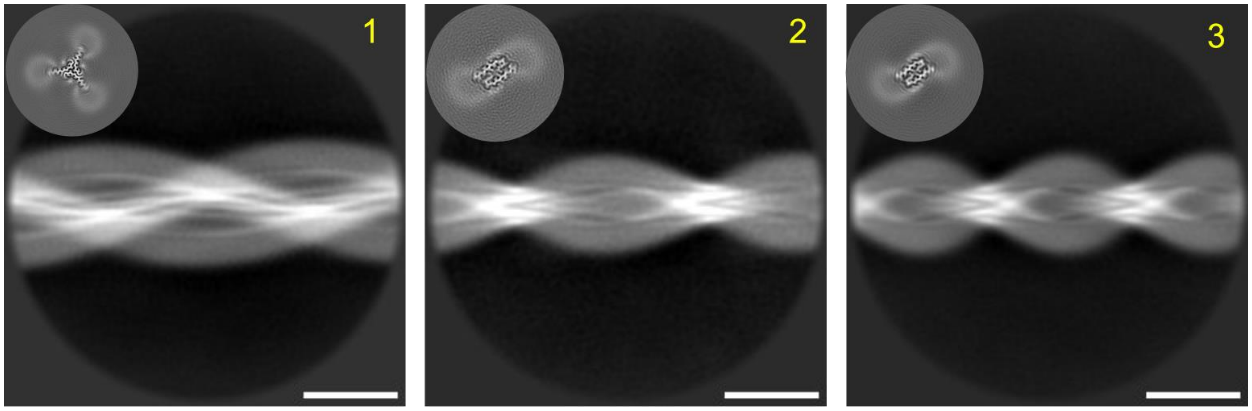


**Figure S8**. **Two-dimensional details of citropin 1.3 amyloids formed with liposomes**. Cryo-EM 2D class averages of different polymorphs obtained in PBS pH 7.4 in the presence of negatively charged liposomes in a 1:5 molar ratio (peptide:liposome). The polymorphs are numbered 1 to 3 representing 24.6%, 24.0%, 12.1% relative particle count per class on total percentage. The insets show the cross-sections of relative class maps. Scale bar 100 Å.

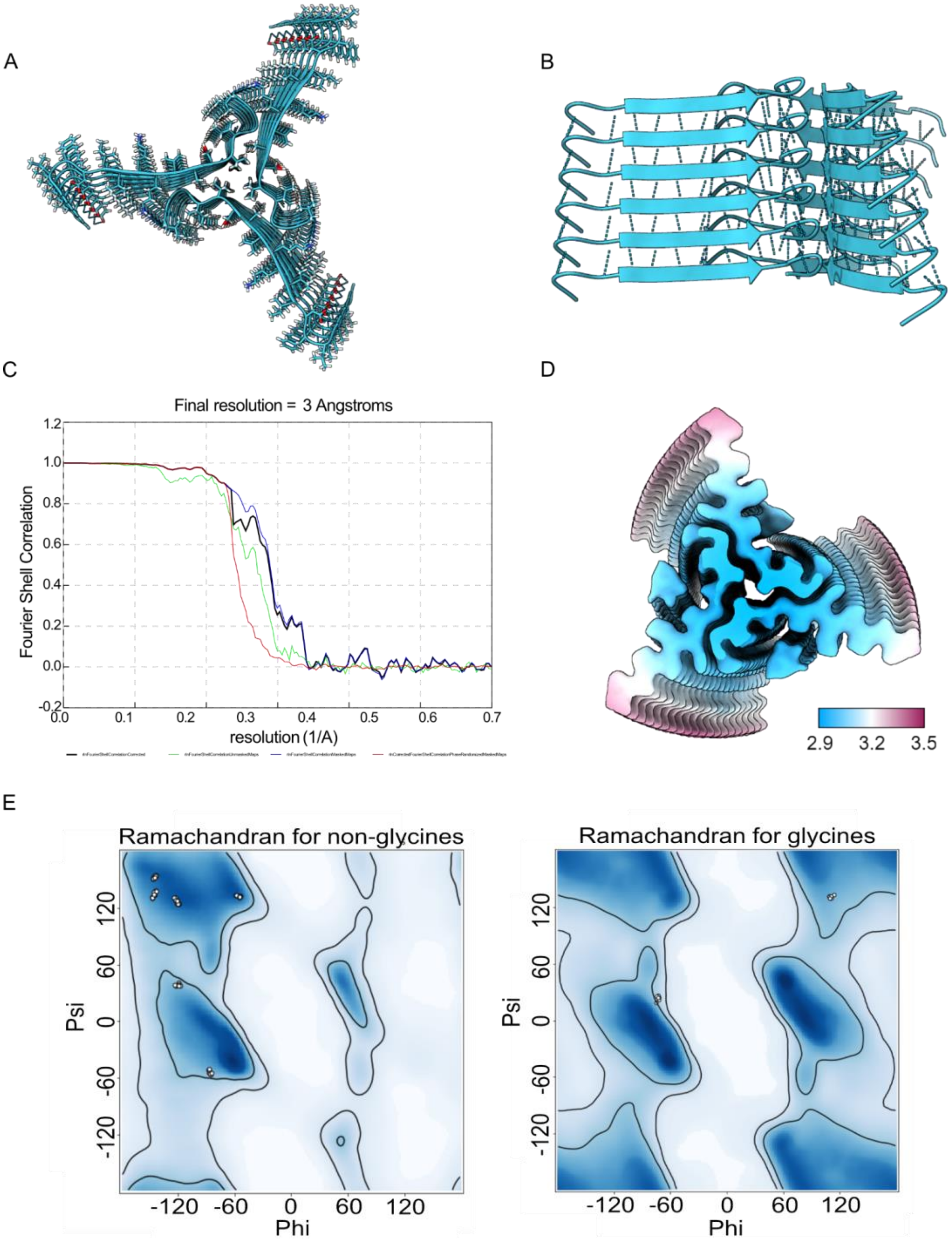


**Figure S9**. **Citropin 1.3 polymorph I-L cryo-EM map and model overviews. A)** A top view of the Pol I-L model with side chains shown as sticks. **B)** Side view with visualization of backbone hydrogen bonds. **C)** GSFC curve reported from Relion of Pol II-L map. **D)** Local resolution visualization of map cross-section. **E)** Ramachandran plots as reported from software Phenix for non-glycine and glycine residues.

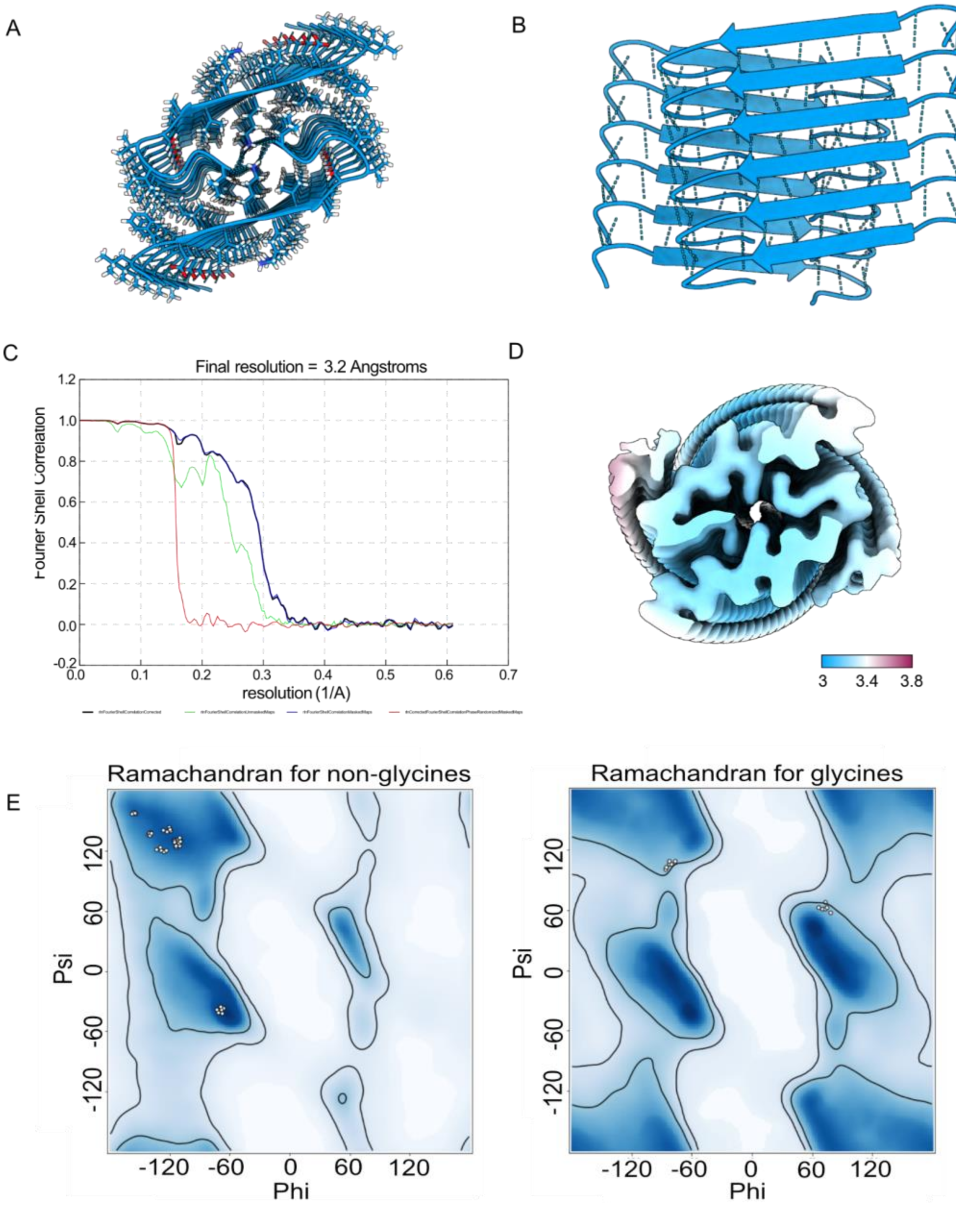


**Figure S10**. **Citropin 1.3 polymorph II-L cryo-EM map and model overviews. A)** A top view of the Pol II-L model with side chains shown as sticks. **B)** Side view with visualization of backbone hydrogen bonds. **C)** GSFC curve reported from Relion of Pol II-L map. **D)** Local resolution visualization of map cross-section. **E)** Ramachandran plots as reported from software Phenix for non-glycine and glycine residues.

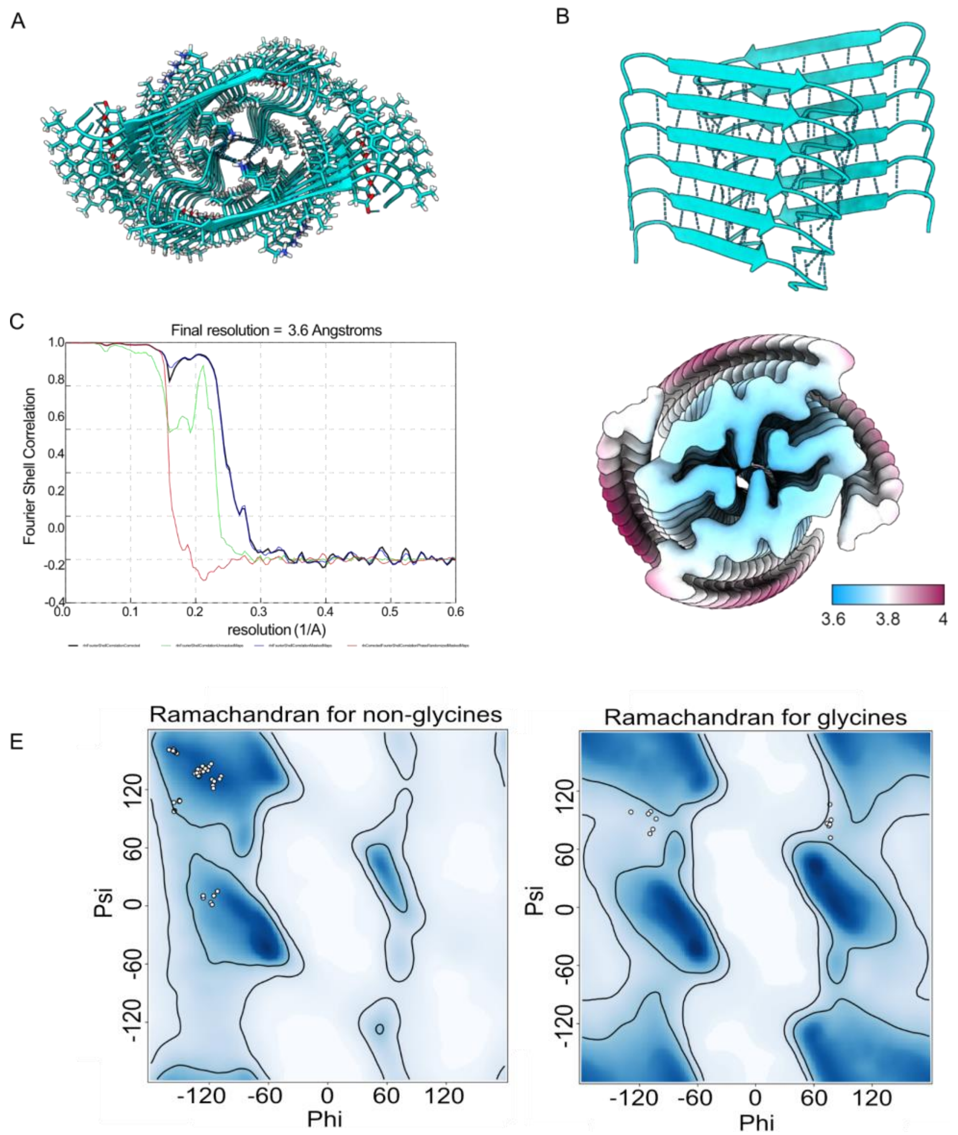


**Figure S11. Citropin 1.3 polymorph III-L cryo-EM map and model overviews. A)** A top view of the Pol III-L model with side chains shown as sticks. **B)** Side view with visualization of backbone hydrogen bonds. **C)** GSFC curve reported from Relion of Pol III-L map. **D)** Local resolution visualization of map cross-section. **E)** Ramachandran plots as reported from software Phenix for non-glycine and glycine residues.

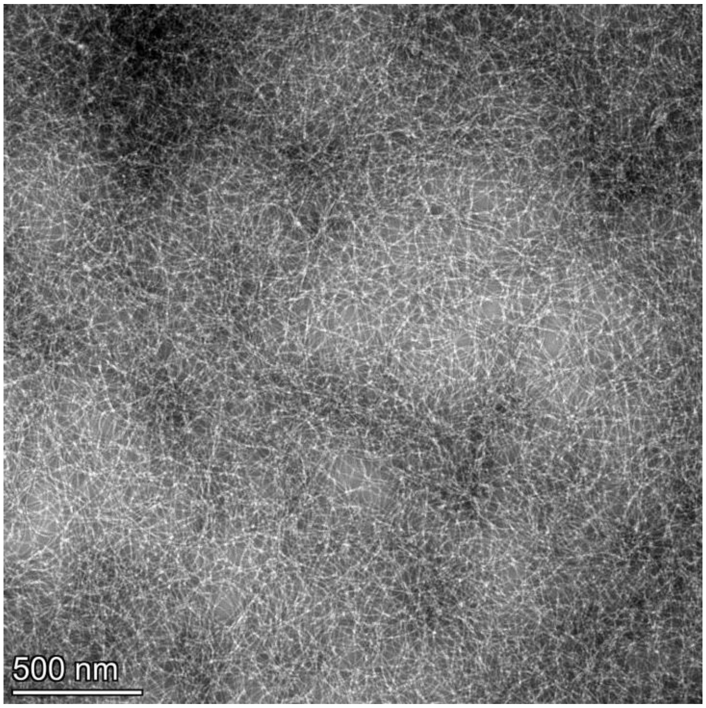


**Figure S12. Negative stain TEM image of FITC-citropin 1.3.** Negative stain TEM image of citropin 1.3 labeled with FITC at the N-terminal end, incubated for 24 hours at 0.5 mM in water. Scale bar 500 nm.

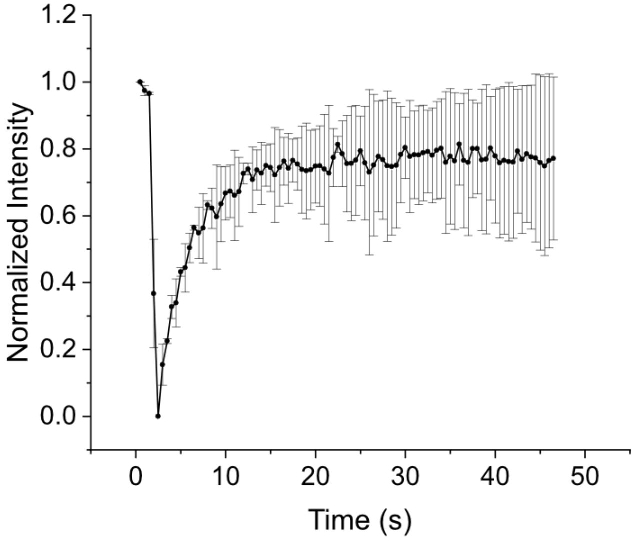


**Figure S13. Fluorescence recovery after photobleaching of droplets.** FRAP results after the interaction between DOPE:DOPG GUVs and citropin 1.3, as visualized in Figure 5. The graph presents an average and error bars based on five measurements from two independent experiments.

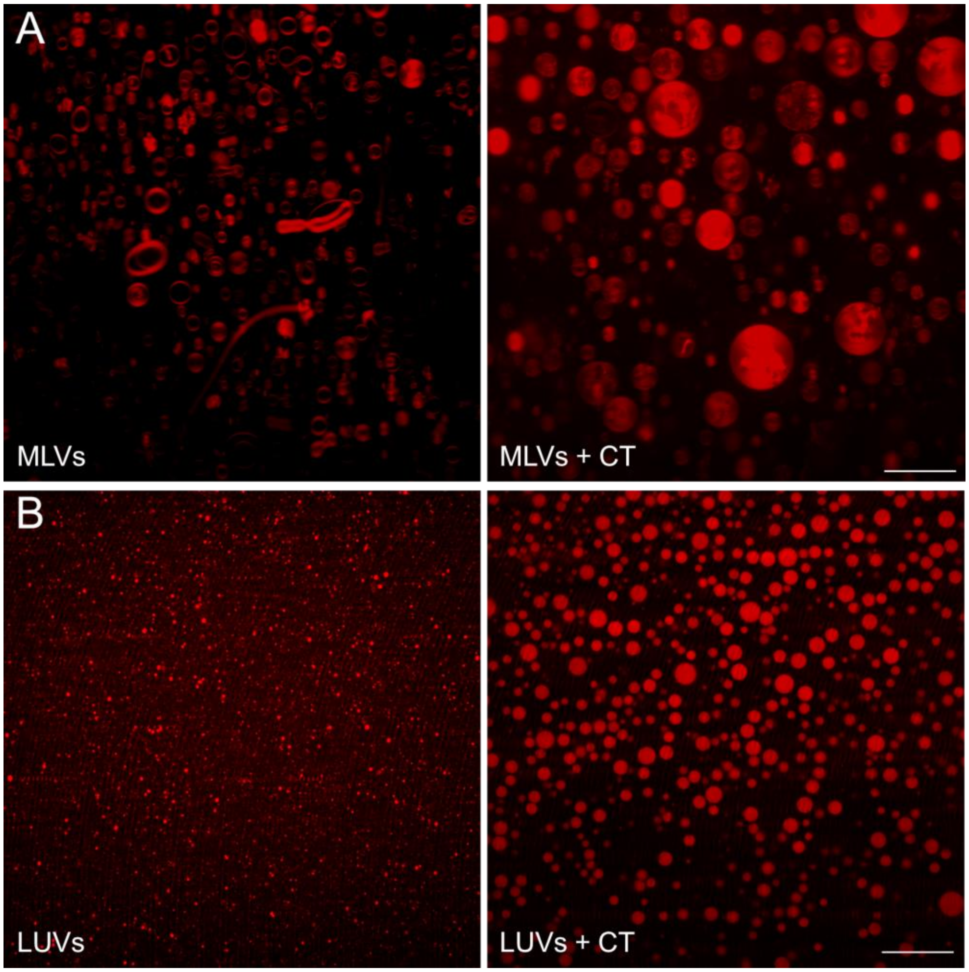


**Figure S14. The effect of citropin 1.3 on liposomes composed of multilamellar and large unilamellar vesicles.** Fluorescence microscopy visualization of: **A)** multilamellar vesicles before (left) and after (right) the addition of citropin 1.3 (CT). **B)** Large unilamellar vesicles before (left) and after (right) the addition of citropin 1.3 (CT). Both panels show the formation of larger vesicles induced by peptide addition. Scale bar 20 μm.

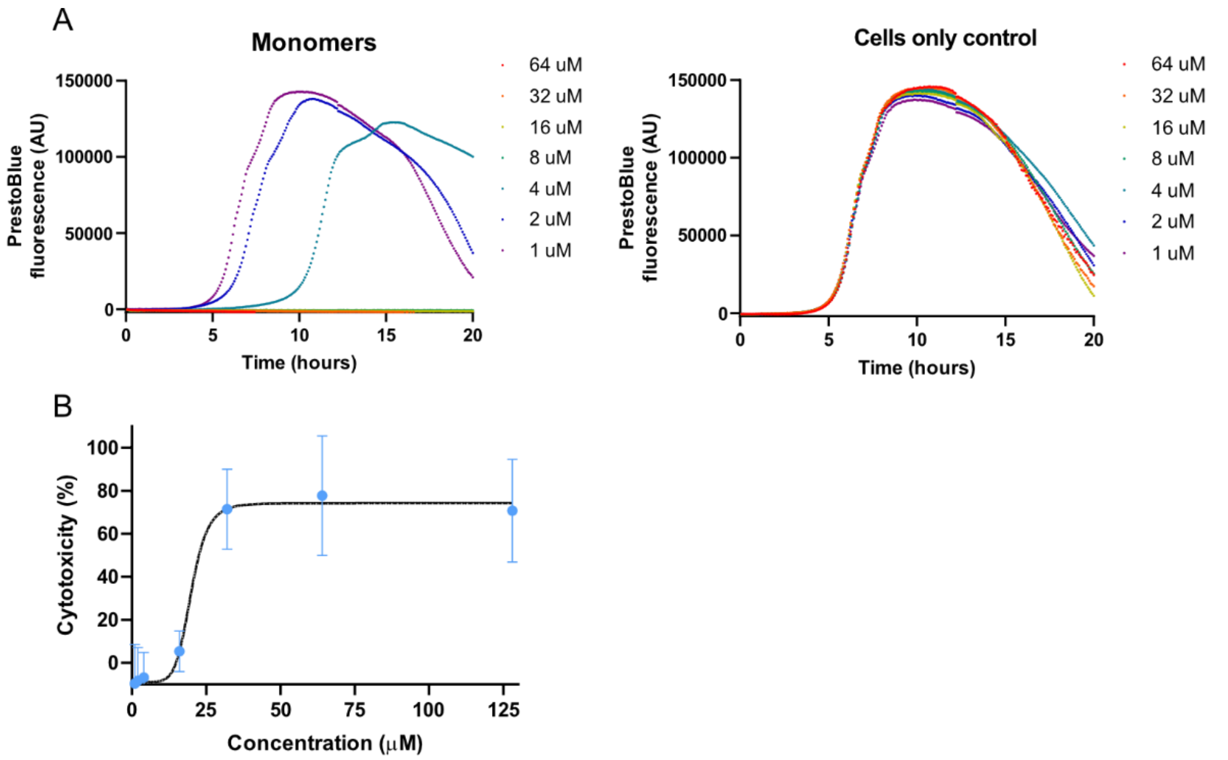


**Figure S15. MIC and $LC_{50}$ values for citropin 1.3. A)** The left panel illustrates the growth of *Bacillus subtilis* in the presence of varying concentrations of citropin 1.3 (1 to 64 μM, as indicated by the color codes). The minimum inhibitory concentration (MIC) was determined to be 8 μM, which was sufficient to completely inhibit bacterial growth. The right panel depicts control experiments, where *Bacillus subtilis* growth was measured with the addition of a buffer at equivalent volumes corresponding to the peptide concentrations (color-coded to match the respective peptide volumes). **B)** The cytotoxicity assay for citropin 1.3 against A-549 cells. The lethal concentration at which 50% of the cells are killed ($LC_{50}$) was determined to be 21 μM.

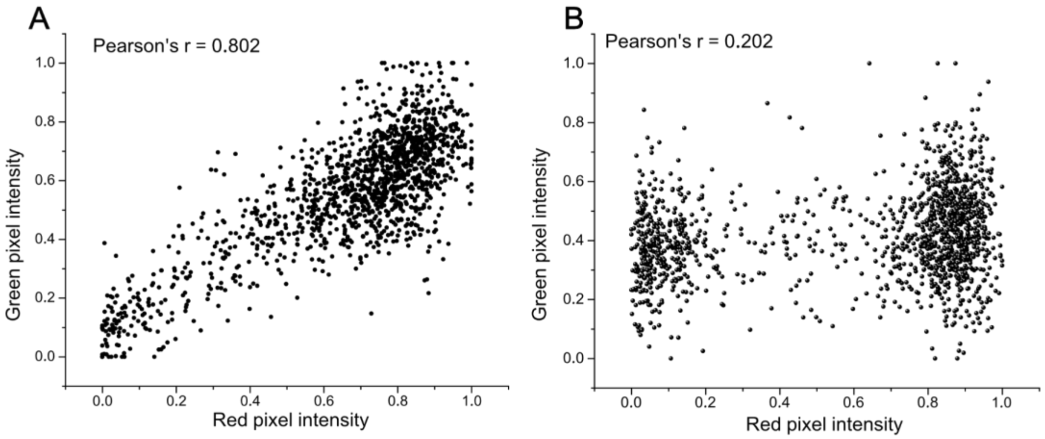


**Figure S16. Scatter plot analysis of normalized red (propidium iodide) and green (FITC-labeled citropin 1.3) pixel intensities within the nucleolus of A-549 cells. A)** Scatter plot showing the distribution of pixel intensities between propidium iodide (red pixel) and 21 μM citropin 1.3 (green pixel, 1% FITC-labeled and 99% unlabeled). **B)** Positive control scatter plot in the absence of citropin 1.3, with Triton X added to induce cell death. The Pearson's r coefficient for each experimental condition is displayed, providing a quantitative measure of the correlation between red and green pixel intensities.

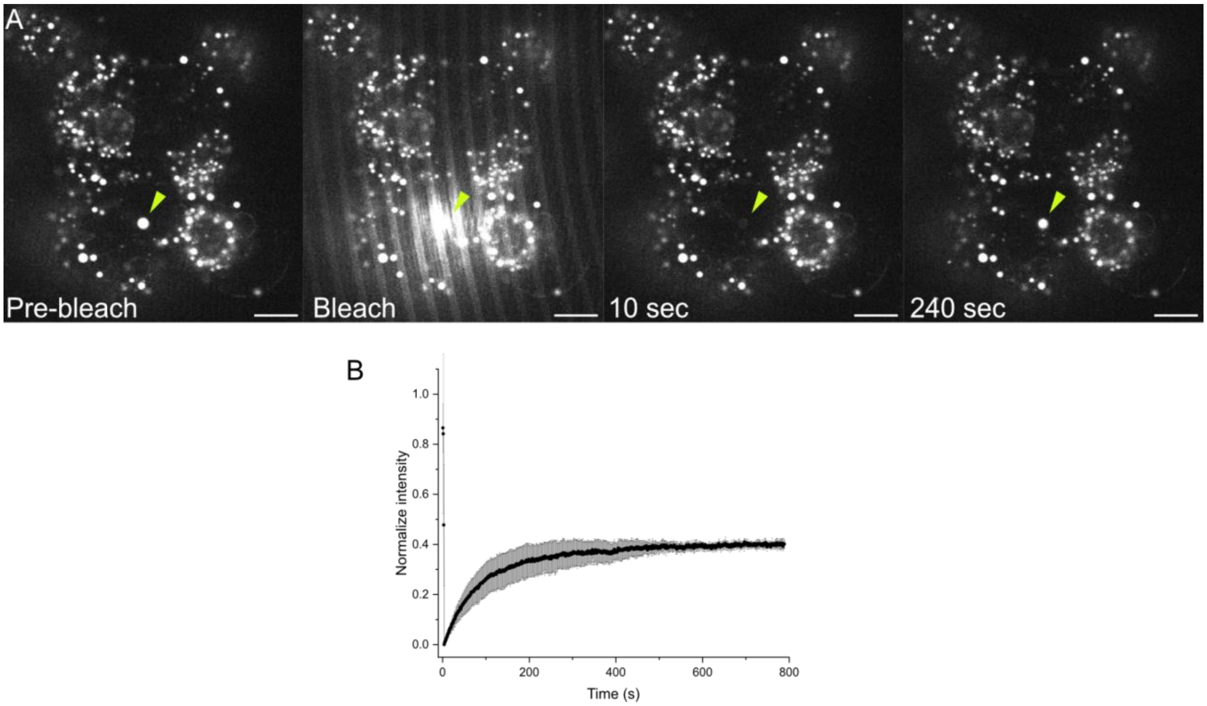


**Figure S17. Fluorescence recovery after photobleaching (FRAP) of droplets formed by citropin 1.3 following A-549 cell lysis**. **A)** FRAP experiment performed on a single droplet enriched with citropin 1.3 (marked by a yellow triangle) demonstrates fluorescence intensity recovery over time after bleaching. **B)** Graphical representation of the FRAP results, showing the average fluorescence recovery curve from four independent measurements. Scale bar 10 µm.

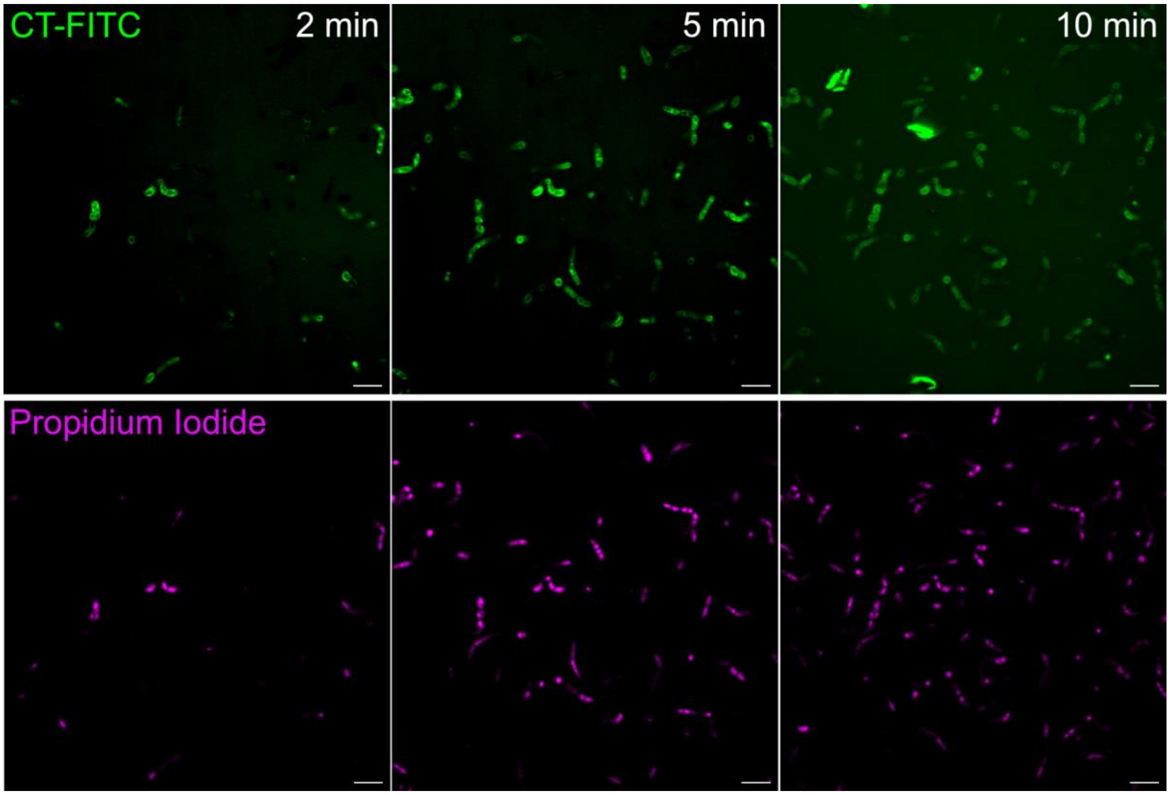


**Figure S18**. **Citropin 1.3 interactions with *Bacillus subtilis*.** Fluorescence microscopy visualization of FITC-labeled citropin 1.3 (CT-FITC) interacting with the bacterial cells, causing cell death as indicated by propidium iodide. Scale bar 5 μM.

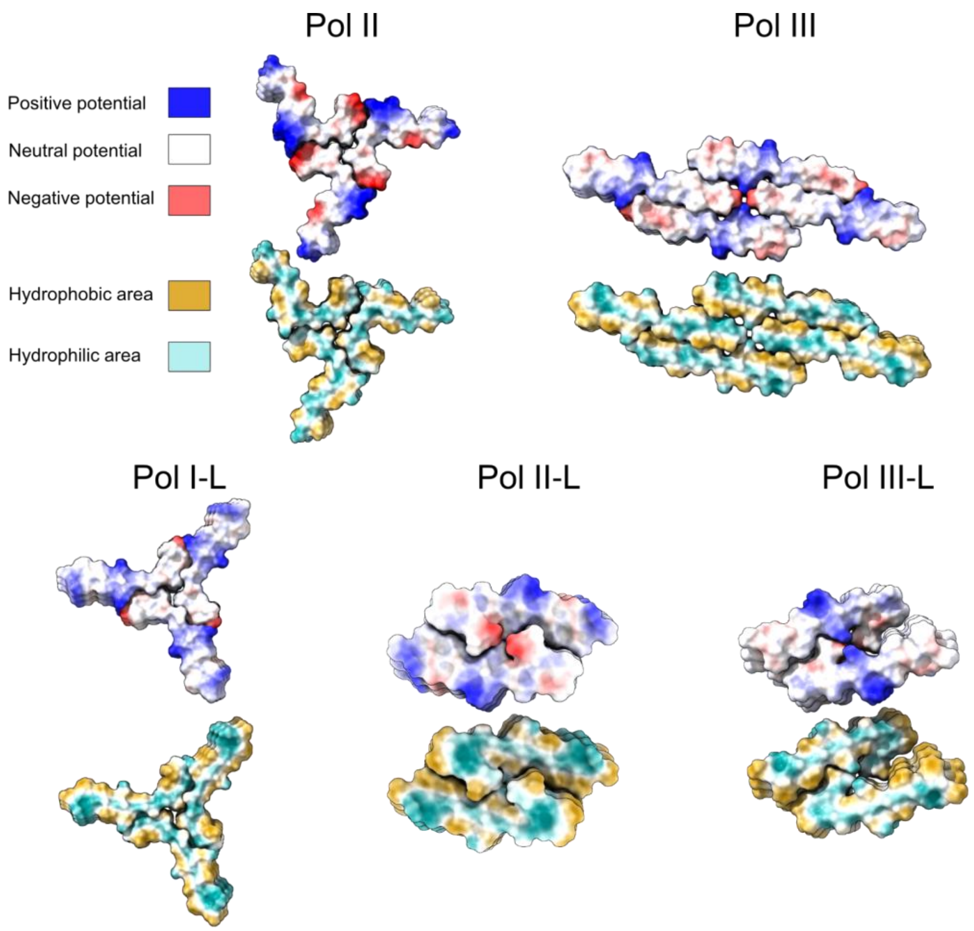


**Figure S19. Electrostatic and hydrophobic surfaces of citropin 1.3 polymorphs.** Surface representation of different citropin 1.3 polymorphs colored by calculated electrostatic and hydrophobic surface. Color codes of surface properties are shown.
